# Opposing Range-Dependent Interactions Create Complex Spatial Patterns of Antibiotic Tolerance in Multispecies Biofilms

**DOI:** 10.64898/2026.02.04.703747

**Authors:** Giulia Bottacin, Benjamin Raach, Leonard Fröhlich, Jasmin Künnecke, Andreas Kaczmarczyk, Alejandro Tejada-Arranz, Giovanni Stefano Ugolini, Roman Stocker, Urs Jenal, Dirk Bumann, Petra S. Dittrich, Olga T. Schubert, Simon van Vliet

**Affiliations:** Biozentrum, University of Basel, 4056 Basel, Switzerland; Department of Environmental Systems Science, ETH Zurich, 8092 Zurich, Switzerland; Department of Environmental Microbiology, Eawag: Swiss Federal Institute of Aquatic Science and Technology, 8600 Dübendorf, Switzerland; Department of Biosystems Science and Engineering (D-BSSE), ETH Zurich, 4056 Basel, Switzerland; Institute of Environmental Engineering, ETH Zurich, 8093 Zurich, Switzerland; Department of Fundamental Microbiology, University of Lausanne, 1015 Lausanne, Switzerland

**Keywords:** multispecies biofilm, spatial structure, interspecies interactions, emergent properties, antibiotic tolerance

## Abstract

Many microbial communities form multispecies biofilms where cells interact through diffusible molecules. In these biofilms, multiple interactions, often with opposing effects, occur simultaneously, yet we lack quantitative frameworks to predict how they combine to shape community functions. Here, we hypothesized that complex spatial patterns can emerge when opposing interactions have distinct spatial ranges. To test this, we studied how two *Pseudomonas aeruginosa* exoproducts, HQNO and rhamnolipids, jointly modulate *Staphylococcus aureus* antibiotic tolerance by respectively increasing and decreasing it. Using microfluidics-based imaging, we quantified spatial-tolerance patterns at single-cell resolution and found that tolerance indeed shows a complex spatial pattern: *S. aureus* cells survived treatment only at intermediate distances from *P. aeruginosa*, while cells closer or farther away did not. Combining experiments and modelling, we showed that this remarkable pattern emerges because rhamnolipids have a stronger but short-ranged effect, while HQNO has a weaker but longer-ranged effect. We found that spatial arrangement affects overall tolerance by shifting the balance between the two opposing interactions. Finally, using bioprinting, we confirmed that HQNO and rhamnolipids modulate tolerance in highly mixed biofilms. In more segregated biofilms, spatial arrangement still strongly modulated tolerance, but independently of these compounds, suggesting additional interactions. Together, our results show that spatial-tolerance patterns emerge from the combined effect of opposing range-dependent interactions and cannot be predicted from either alone. By predicting how opposing interactions jointly determine community properties, our framework provides a foundation for understanding and ultimately engineering microbiome functions.

## Introduction

Biological systems often display properties that cannot be deduced from their parts alone, as these properties emerge from how these parts interact and organize in space (1). Turing patterns provide a classic example of how multiple interactions can combine to form complex spatial patterns (2). They emerge from the combined effect of two interactions with opposing signs and different ranges, and are thought to underlie morphogen-driven patterning in nature, from the stripes on zebrafish (3, 4) to the arrangement of feathers (5) and limbs (6). They have also been linked to species distributions of plants (7, 8), gene expression patterns in synthetic microbial systems (9), and the regular spacing of heterocysts in filamentous cyanobacteria (10). While Turing patterns are a well-known example, the underlying principle is more general: complex spatial patterns can emerge whenever multiple interactions with different signs and ranges act on spatially structured systems. Here, we investigate how such processes affect multispecies biofilms.

Microbial communities often form dense, surface-attached assemblies such as biofilms, where high cell density and limited mobility cause cells to become spatially organized (11). The properties of these biofilms emerge from complex networks of molecular interactions in which cells both positively and negatively affect each other’s growth, activity, and survival (12, 13). Most of these interactions are mediated through the exchange of diffusible compounds, with interaction strength decaying with distance between cells (14–18). The spatial range of each interaction is determined by cell densities and by the uptake and diffusion rates of the exchanged compounds, and thus differs between interactions (17, 18). Consequently, both the microscale spatial organization of cells and the molecular mechanisms underlying interactions determine which cells can interact with each other (18).

Previous studies have characterized how spatial structure affects communities by examining individual interaction types in isolation. For example, spatial segregation can stabilize intraspecies cooperation and protect cells from antagonistic interactions (19, 20), but can also hinder mutualistic interspecies cross-feeding (21–23). While the spatial effects of such isolated interactions are relatively well understood, natural communities involve complex interaction networks where multiple interactions of different signs, ranges, and strengths act simultaneously. However, we lack quantitative frameworks to predict how such interactions combine to shape emergent properties in multispecies biofilms.

To investigate the combined effect of multiple opposing interactions, we used a model community of *Pseudomonas aeruginosa* and *Staphylococcus aureus*, focusing on how *P. aeruginosa* modulates *S. aureus* antibiotic tolerance. *P. aeruginosa* produces multiple exoproducts that affect *S. aureus* tolerance to tobramycin (24–27). We focused on two well-characterized molecules with opposing effects (Fig. 1A): 2-Heptyl-4-Quinoline N-Oxide (HQNO) and rhamnolipids. HQNO increases tolerance by inhibiting cytochrome oxidases, collapsing the proton motive force (PMF) required for aminoglycoside uptake (24, 28). Rhamnolipids decrease tolerance by increasing membrane permeability, which enhances PMF-independent aminoglycoside uptake (24, 25).

**Fig. 1.**
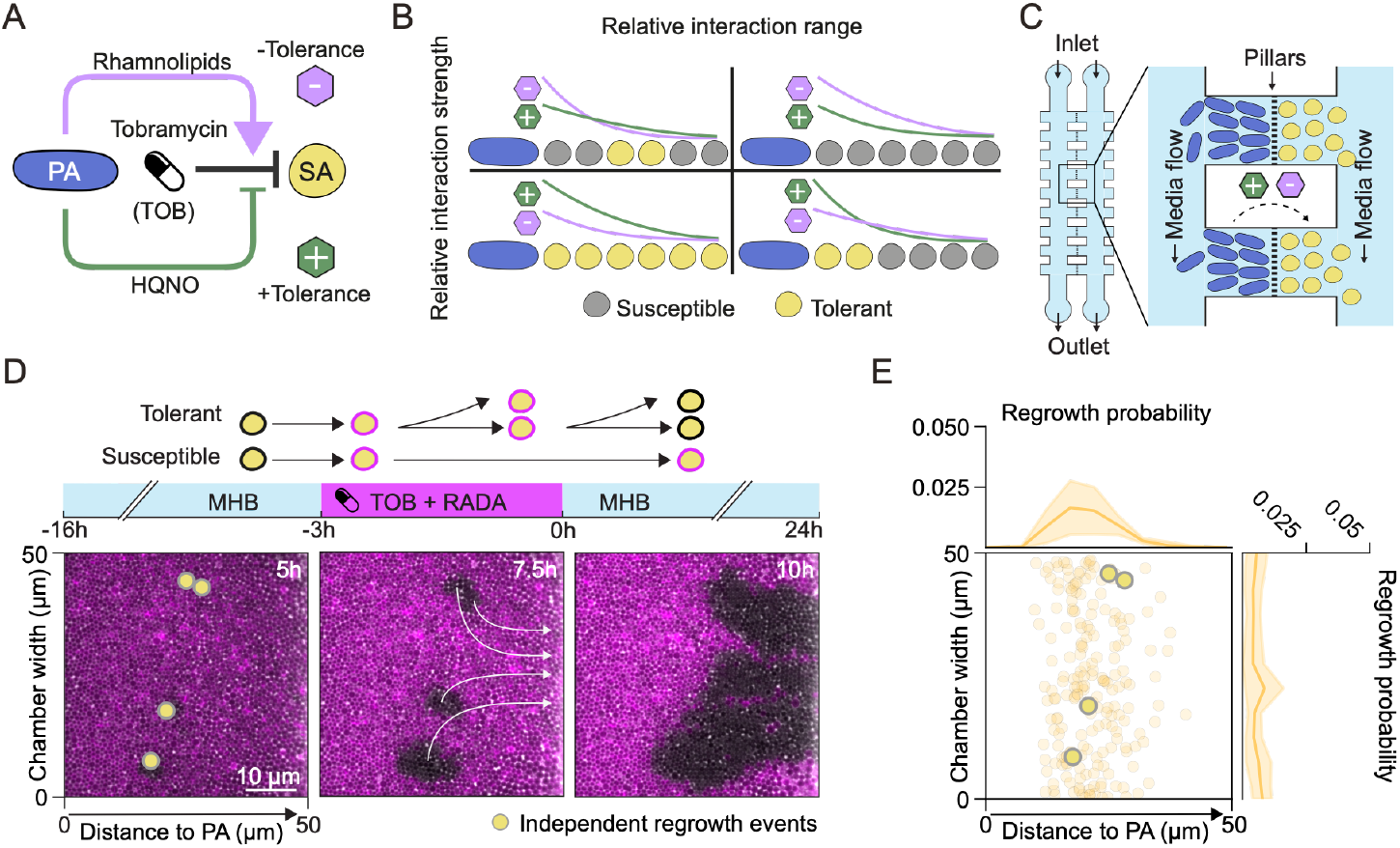
Antibiotic tolerance in *S. aureus* peaks at intermediate distances from *P. aeruginosa*. (A) *P. aeruginosa*-secreted rhamnolipids and HQNO have opposing effects on *S. aureus* tolerance to tobramycin (TOB): Rhamnolipids decrease tolerance by facilitating TOB uptake, while HQNO increases tolerance by reducing uptake. (B) In spatially structured environments, HQNO and rhamnolipids form gradients. Depending on each compound’s relative range and strength, four distinct spatial tolerance patterns are possible. (C) Microfluidic device with paired chambers (50 × 50 × 0.8 μm) separated by a pillar array that permits compound diffusion but maintains species segregation. Each chamber contains ∼1000 cells in a monolayer. Flow channels on either side of the paired chambers provide continuous nutrient flow and remove excess cells. (D) Top: Tolerance assay protocol with 3 hours TOB treatment (58 μg/ml, 64 × MIC) followed by 24 hours recovery. During treatment, *S. aureus* cells were stained with RADA (fluorescent D-amino acid); after treatment, growing cells dilute the dye while non-growing cells remain stained. Bottom: Time series of an *S. aureus* chamber at 5, 7.5, and 10 hours post-treatment showing phase contrast (grey) and RADA fluorescence (magenta). Regrowing cells appear black; non-growing cells remain magenta. Clusters of regrowing cells push surrounding cells toward the flow channel (white arrows). Yellow circles mark locations of independent regrowth events. Images were cropped and adjusted equally for brightness and contrast. (E) Spatial distribution of independent regrowth events across distance from *P. aeruginosa* and chamber width. Large dots: chamber from panel D; small dots: all regrowth events from 29 chambers across 3 independent experiments (n=10 per experiment).

While the molecular mechanisms underlying these interactions have been extensively characterized, how they combine to influence community-level outcomes in a spatial context remains unclear. Previous work in a mouse chronic wound model demonstrated that spatial structure enhances *S. aureus* tolerance to tobramycin, and that this protective effect is eliminated by homogenization or by loss of HQNO production, linking the spatial dependence of tolerance to HQNO (29). However, this study did not consider the effect of rhamnolipids or other *P. aeruginosa* products that can affect tolerance in *S. aureus*. Because these molecules differ in their effects and likely their ranges, how they combine to determine tolerance in spatially structured communities remains unknown.

We hypothesized that complex spatial patterns can emerge when multiple interactions with opposing effects and different ranges target the same phenotype. Because interaction strengths decay at different spatial scales, their net effect varies with distance from the producer, creating spatial variation in recipient phenotypes. Specifically, we expect *S. aureus* tolerance to vary with distance from *P. aeruginosa*. Depending on the relative strength and range of the two opposing interactions, different tolerance patterns would emerge, ranging from localized zones of protection to overall susceptibility (Fig. 1B).

To test our hypothesis, we grew the community in a microfluidic device to precisely control spatial arrangement and measure growth and tolerance at single-cell resolution. We found that *S. aureus* survival was highest at intermediate distances from *P. aeruginosa*. By combining experimental analyses of mutant strains deficient in HQNO and/or rhamnolipid production with a reaction-diffusion model, we showed that this pattern results from rhamnolipids having a shorter interaction range but higher relative strength near *P. aeruginosa*, while HQNO has a longer range that dominates at intermediate distances. Finally, using experiments with microfluidic chambers and bioprinted colony biofilms, we demonstrated that microscale spatial arrangement is an important determinant of overall tolerance levels in *S. aureus*.

## Results

### Antibiotic tolerance peaks at intermediate distances from *P. aeruginosa*

To quantify how multiple interactions jointly shape community-level tolerance, we used a microfluidic device in which *P. aeruginosa* and *S. aureus* grow as monolayers in adjacent chambers connected by a porous wall consisting of closely spaced pillars (Fig. 1C). The gaps between pillars allow for the exchange of diffusible compounds while preventing physical contact between cells. The chambers open into two independent flow channels which supply nutrients and flush out excess cells.

We first confirmed that HQNO and rhamnolipids are produced in our system and modulate *S. aureus*’s tolerance to tobramycin. Using batch-culture supernatant assays, we validated previous findings (24) showing that HQNO increases tolerance while rhamnolipids decrease it (Fig. S1). Using transcriptional reporters, we further demonstrated that *P. aeruginosa* produces both compounds in the microfluidic system (Fig. S2).

To test how *P. aeruginosa* affects tolerance in *S. aureus*, we co-cultured the two species overnight in Mueller Hinton Broth (MHB), followed by 3 hours of treatment with the antibiotic tobramycin at 64 x MIC (58 μg/ml). To distinguish between cells that could and could not regrow after treatment, we added red fluorescently labeled D-Amino Acids (RADA) that stain the cell’s peptidoglycan during the treatment phase. After treatment, the medium was switched back to MHB for 24 hours. Tolerant *S. aureus* cells capable of regrowth dilute the RADA, and thus lose fluorescence, whereas non-regrowing susceptible cells remain stained (Fig. 1D, SI Movie 1).

In the absence of *P. aeruginosa*, we observed no regrowth of *S. aureus* cells after antibiotic treatment in most chambers (65 out of 68) (Fig. S3, SI Movie 2). However, in the presence of *P. aeruginosa*, we found that in most chambers (26 out of 29) a small fraction of *S. aureus* regrew (Fig. S3).

To quantify how proximity to *P. aeruginosa* shaped *S. aureus* antibiotic tolerance, we developed an automated image analysis pipeline that identifies the location of independent regrowth events (Fig. S4). As these first regrowing cells divide, they give rise to increasingly large clusters that push surrounding cells towards the flow channel (Fig. 1D). Because of subsequent cluster merging and cell loss to the flow channel, our analysis likely underestimates the absolute number of regrowth events. However, such underestimation would only affect our conclusions if it were spatially biased. To test this, we compared distributions of regrowth locations in the middle of the chamber (where cells move rapidly) and at the edges (where movement is slower). We observed similar distributions in both regions (Fig. 1E), indicating that while we may miss some regrowth events, this does not systematically bias the spatial patterns we observe.

By mapping the spatial distributions of surviving *S. aureus* cells, we found a non-monotonic dependence of survival on distance, with survival peaking at intermediate distances from *P. aeruginosa* (Fig. 1E). Survival showed significant non-uniformity with distance to *P. aeruginosa* (χ^2^ = 242.75, df = 9, p < 0.001). In contrast, survival along the chamber width showed no significant deviation from uniformity (χ^2^ = 14.98, df = 9, p = 0.059). Together, these results indicate that survival depends specifically on distance from *P. aeruginosa*.

While initial regrowth events were confined to an intermediate spatial range from *P. aeruginosa*, even within this zone only a small fraction of cells (2%) resumed growth. This indicates that the chemical gradients determine the spatial region where tolerance is possible but that regrowth within this region remains a stochastic event at the single-cell level.

### Spatial tolerance patterns emerge from opposing effects of HQNO and rhamnolipids

To determine how HQNO and rhamnolipids shape the observed tolerance patterns, we repeated the microfluidic tolerance assays using *P. aeruginosa* mutants deficient in the production of either or both compounds. Co-culture with the *ΔrhlA* mutant, which cannot produce rhamnolipids, led to a 4-fold increase in overall survival (Fig. 2A, SI Movie 3). This increase was not uniform in space (Fig. 2B). Rather, *S. aureus* survival increased by 145-fold in close proximity to *P. aeruginosa* but by only 3.8-fold at intermediate distances (Fig. S5), indicating that rhamnolipids have a short-range tolerance-reducing effect. Loss of rhamnolipids also decreased the recovery time needed for cells to resume growth (Fig. S6). In contrast, co-culture with the *ΔpqsL* mutant, which cannot synthesize HQNO, resulted in uniformly low *S. aureus* survival, with only four regrowth events observed across all 30 chambers (Fig. 2). The *ΔpqsLΔrhlA* double mutant showed similarly low survival (Fig. 2), indicating that, under our conditions, HQNO is necessary to promote tolerance.

**Fig. 2.**
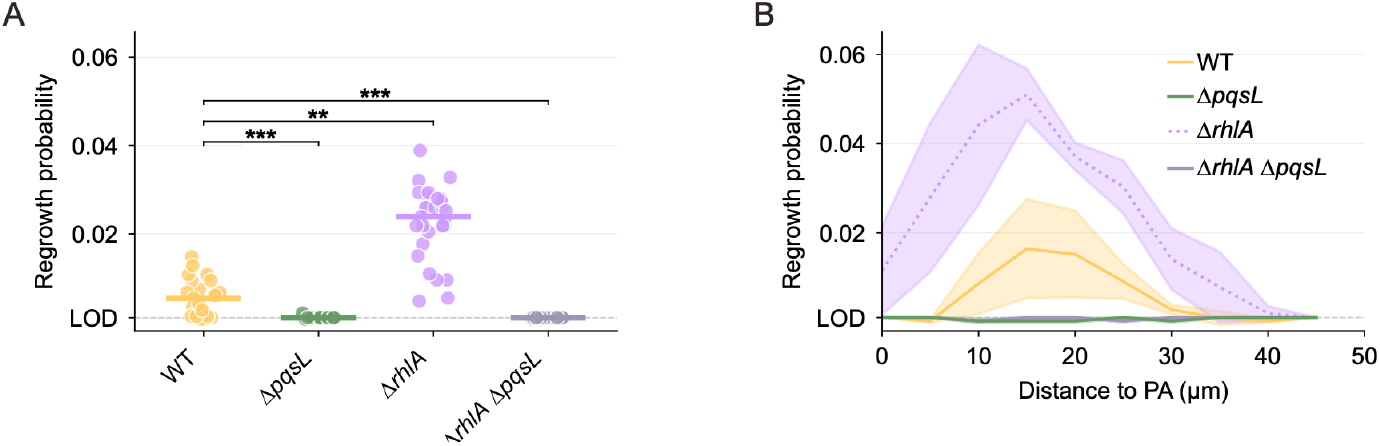
Spatial tolerance emerges from combined effect of HQNO and rhamnolipids. **(A)** Average regrowth probability of *S. aureus* co-cultured with WT *P. aeruginosa* (yellow) or mutants not able to produce HQNO (Δ*pqsL*, green), rhamnolipids (Δ*rhlA*, purple), or both (Δ*rhlA* Δ*pqsL*, grey). Points: mean regrowth fraction in a single chamber; lines: median across all chambers. Values below the limit of detection (LOD = 0.001) are shown at the LOD. Statistical differences were assessed using Kruskal-Wallis test followed by Dunn’s post-hoc test with Bonferroni correction for pairwise comparisons against WT. *p < 0.05, **p < 0.01, ***p < 0.001. **(B)** Regrowth probability of *S. aureus* as a function of distance from *P. aeruginosa* for WT (yellow, same data as Fig. 1E), Δ*pqsL* (green), Δ*rhlA* (purple), or Δ*pqsL* Δ*rhlA* (grey). Lines: mean regrowth fraction per spatial bin (5 µm); shaded areas: ± standard deviation across chambers. Data for each strain from 29 chambers (WT or *ΔrhlA*) or 30 chambers (*ΔpqsL* or *ΔrhlA ΔpqsL*) across 3 independent devices (n=10 per device)

Although the rhamnolipid-deficient mutant (*ΔrhlA*) extended the zone of protection towards the interface, *S. aureus* tolerance remained lower close to *P. aeruginosa* than at intermediate distances. We performed several control experiments to rule out potential experimental artifacts that could cause this pattern. First, we examined whether the spatial tolerance pattern could result from differences in pre-treatment growth rate. We measured *S. aureus* growth rates as a function of distance from *P. aeruginosa* before treatment and found no significant spatial variation, ruling out growth rate as a cause of the observed pattern (Fig. S7). Second, we tested whether lysis products released by *P. aeruginosa* during treatment could affect *S. aureus* tolerance. Flushing *P. aeruginosa* out of the chambers before treatment did not change the survival pattern, indicating that the pattern emerges from interactions prior to treatment (Fig. S8). Third, we tested whether spatial gradients in antibiotic concentration affect tolerance patterns by supplying antibiotics only from the flow channel on the *S. aureus* side, ensuring that antibiotic concentrations would be lowest near the porous wall. We observed no change in the survival patterns, ruling out potential antibiotic gradients as an explanation (Fig. S9).

Another explanation for the lower survival in *S. aureus* close to *P. aeruginosa* is that *P. aeruginosa* produces other short-ranged compounds that decrease tobramycin tolerance in *S. aureus*, beyond rhamnolipids. While rhamnolipids are the only known compound secreted by *P. aeruginosa* that decreases tolerance to aminoglycosides, several other compounds can decrease tolerance to other antibiotics. For example, LasA endopeptidase decreases tolerance to vancomycin (24) while multiple LasR-regulated factors decrease tolerance to norfloxacin (30).

To explore which factor might mediate the additional tolerance-reducing interaction, we performed an initial batch culture screen. Using a previously constructed *P. aeruginosa* two-allele transposon mutant library (31), we selected mutants deficient in effectors previously linked to tolerance modulation and exposed *S. aureus* to their supernatants prior to tobramycin treatment. However, these experiments did not reveal any single-gene disruptions that significantly decreased *S. aureus* tolerance (Fig. S10). Our screen tested only a subset of candidate genes, and differences in culture conditions and strain backgrounds between the batch screen and microfluidic experiments could affect which interactions are detectable. An additional short-ranged tolerance-reducing interaction therefore remains the most likely explanation for the observed pattern. We leave identification of this putative interaction to future work.

Taken together, our results suggest that the spatial tolerance pattern emerges from opposing, spatially overlapping effects. At short range, rhamnolipids reduce tolerance, while at longer ranges HQNO dominates, increasing tolerance.

### Fluorescent reporters reveal HQNO and rhamnolipid spatial gradients

To quantify the interaction ranges and strengths, we used a two-step approach. We first estimated spatial concentration gradients using fluorescent reporters. We then used a reaction-diffusion model to infer the interaction parameters. To estimate HQNO and rhamnolipid concentrations, we used fluorescent reporters that respond to the phenotypic changes induced by these molecules (32). HQNO inhibits *S. aureus* respiration, pushing cells toward fermentation (32). As a proxy for this response, we measured the expression of the fermentation-specific gene formate acetyltransferase (*pflB*) using a fluorescent transcriptional reporter (32–34). Rhamnolipids, on the other hand, disrupt the *S. aureus* cell membrane. Following Gdaniec et al. (35), we quantified this disruption using propidium iodide (PI), a dye that enters cells only through damaged membranes and fluoresces upon DNA binding. We confirmed using flow cytometry that both these reporters showed a dose-dependent response to the addition of purified HQNO and rhamnolipids, respectively (Fig. S12B, D).

We then measured the fluorescent response of *S. aureus* grown in co-culture with *P. aeruginosa* in the microfluidic chambers. The fluorescent signal for PI decreased monotonically with distance from *P. aeruginosa* (Fig. S11D). *pflB* expression also decreased with distance beyond approximately 7.5 μm from the interface. However, expression was unexpectedly lower in close proximity to *P. aeruginosa* (Fig. S11A), suggesting interference from additional factors at short range. We therefore fitted the model only on the region beyond 7.5 μm from *P. aeruginosa*.

Finally, we estimated HQNO and rhamnolipid concentrations as a function of distance from *P. aeruginosa* by converting fluorescence intensities into concentrations. We first confirmed that fluorescent intensities scale linearly between flow cytometry and microscopy (Fig. S12A, C). We then fitted dose-response curves to flow cytometry data (Fig. S12B, D), thereby obtaining estimates of local compound concentrations from the observed fluorescence gradients in the microfluidic chambers (Fig. S11B, E).

### Reaction-diffusion modeling reveals distinct ranges and strengths of opposing interactions

To infer the interaction ranges of HQNO and rhamnolipids, we fitted a reaction-diffusion model to the estimated concentration gradients. The model describes the spatial distribution of these two compounds, assuming a constant rate of production by *P. aeruginosa* and a constant rate of uptake by *S. aureus*, while accounting for the diffusive dynamics in the microfluidic setup. The resulting steady state concentration profiles showed that both HQNO and rhamnolipids form gradients that decay with distance from *P. aeruginosa*, each with distinct decay rates (Fig. 3B). The model fit confirmed that rhamnolipids have a shorter interaction range (13.8 μm, 25–75% confidence interval = 12.0–14.3 μm) than HQNO (32.8 μm, 25–75% confidence interval = 26.7–72.5 μm) (Fig. S14A).

**Fig. 3.**
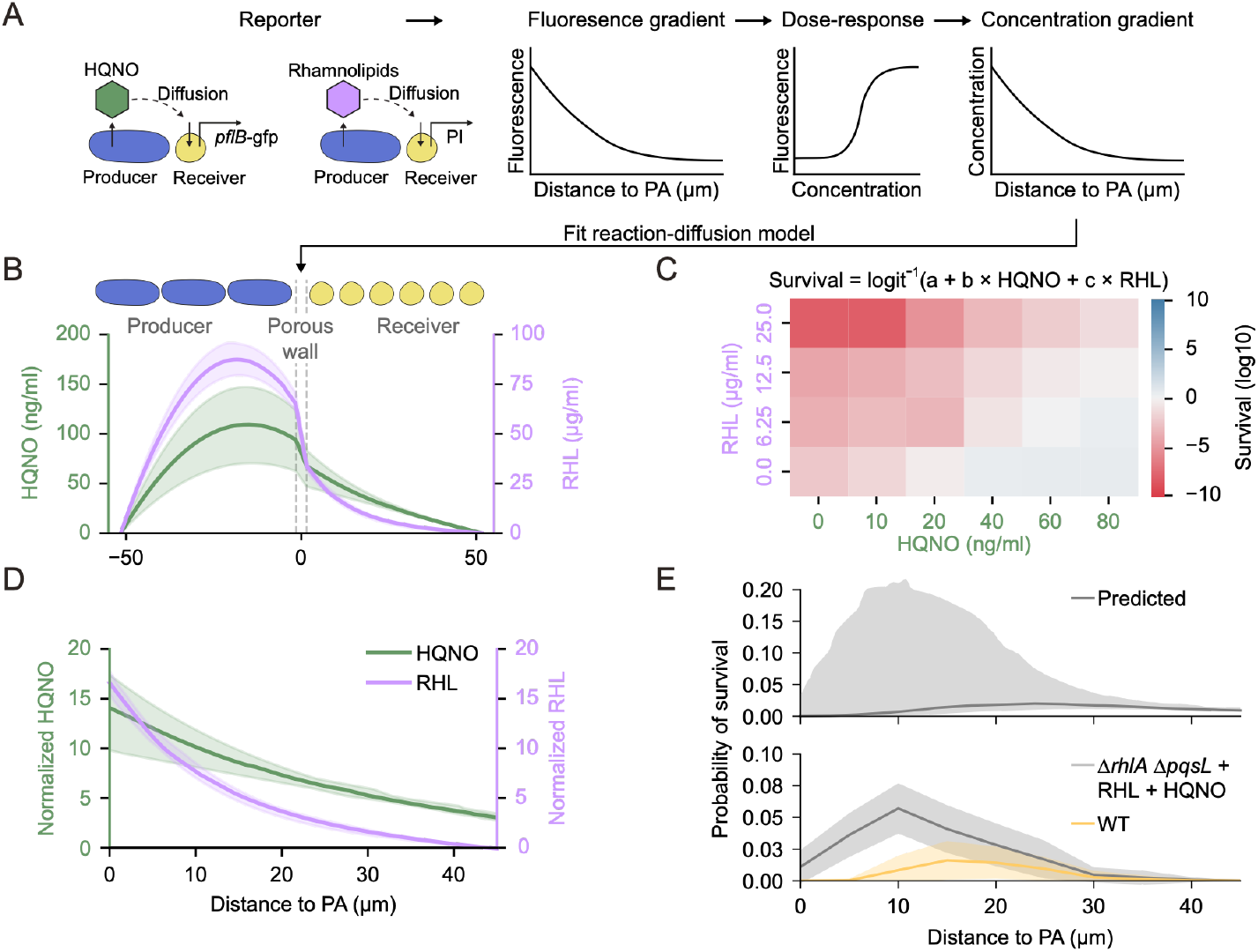
Reaction-diffusion model links molecular gradients to predicted *S. aureus* survival. **(A)** Workflow for estimating local HQNO and rhamnolipid concentrations. Fluorescent reporters (*pflB-gfp* for HQNO, PI for rhamnolipids) were measured as a function of distance from *P. aeruginosa*. Flow cytometry dose-response curves enabled conversion of fluorescence to concentration gradients, which were fitted with a reaction-diffusion model. **(B)** Concentration profiles of HQNO (green, left axis) and rhamnolipids (purple, right axis) versus distance from the porous wall, showing analytical solutions of the reaction-diffusion model with parameters estimated from the data. Lines: median; shaded areas: interquartile range (IQR). **(C)** *S. aureus* survival fraction (CFU/CFU_0_) at 3 hours post-tobramycin exposure (58 µg/mL) across HQNO (ng/mL) and rhamnolipid (µg/mL) concentrations, shown on log_10_ scale. A logistic regression model quantified each compound’s contribution to survival: Survival = logit^−1^ (a + b × HQNO + c × RHL). **(D)** HQNO (green, left axis) and rhamnolipid (purple, right axis) concentrations from panel B scaled by their fitted effect sizes from the logistic model in panel C, enabling direct comparison of each compound’s contribution to survival. Lines: analytical solutions with median parameter estimates; shaded areas: IQR. **(E) Top:** Predicted survival probability from applying the logistic model to fitted concentration profiles. Line: median; shaded area: IQR obtained by exhaustive sampling across parameter estimates. **Bottom:** Observed survival probability versus distance from *P. aeruginosa* for a co-culture of *S. aureus* with wild-type *P. aeruginosa* (yellow, same data as Fig. 1E) and a co-culture with the *P. aeruginosa ΔpqsLΔrhlA* double mutant, supplemented with exogenous HQNO and rhamnolipids (grey). Lines: mean per 5 µm bin; shaded areas: ± standard deviation across chambers.

Having estimated the spatial concentration gradients of HQNO and rhamnolipids, we next quantified the strength of each compound’s effect on antibiotic tolerance. We measured *S. aureus* survival in batch culture across a range of HQNO and rhamnolipid concentrations following tobramycin treatment. Survival increased with HQNO concentration and decreased with rhamnolipid concentration (Fig. 3C). We fitted a logistic survival function to these data, assuming additive effects of the two compounds (Fig. S13). Using the fitted effect sizes, we then scaled the estimated spatial concentrations to directly compare the relative contributions of each compound to survival (Fig. 3D). This analysis revealed that at short distances from *P. aeruginosa*, the tolerance-decreasing effect of rhamnolipids dominates, while at greater distances, the tolerance-increasing effect of HQNO dominates (Fig. 3D).

### Model predictions recapitulate observed spatial tolerance patterns

To validate the model, we predicted *S. aureus* survival probability as a function of distance from *P. aeruginosa* based on the fitted parameters (Fig. 3E, top). The model reproduces the experimentally observed spatial tolerance pattern, with peak survival at intermediate distances, and predicts comparable survival probabilities (Fig. 3E, bottom). However, the model predicts higher survival toward the chamber exit than observed, which can be explained by a limitation of the fluorescent reporters. As cells are pushed toward the exit, they experience decreasing HQNO and rhamnolipid concentrations, but their fluorescence intensities decrease more slowly because GFP and PI must be diluted through growth. This causes us to overestimate concentrations toward the chamber exit, accounting for the observed discrepancy.

To further test the model, we performed a complementation assay with the *P. aeruginosa ΔpqsLΔrhlA* double mutant. We used the model to predict the external HQNO and rhamnolipid concentrations needed to replicate wild-type-like gradients in the double mutant (Fig. S14B) and added them to the *P. aeruginosa* flow channel. The resulting tolerance pattern qualitatively matched that of wild-type *P. aeruginosa*, with highest survival at intermediate distances (Fig. 3E, bottom). However, overall survival probabilities were substantially higher, suggesting we overestimated the HQNO concentration. This discrepancy likely stems from uncertainties in parameter estimation or from overestimating the diffusive blockage caused by the porous wall. Nonetheless, these results demonstrate that the observed tolerance pattern can be explained by the combined effects of two opposing gradients: a strong but short-ranged tolerance-decreasing effect of rhamnolipids and a weaker but longer-ranged tolerance-increasing effect of HQNO.

### Spatial arrangement modulates community-level tolerance

Having established how opposing molecular gradients shape *S. aureus* tolerance in a microfluidic system with fixed spatial arrangement, we next asked how these findings extend to more complex arrangements. We hypothesized that if interactions have different ranges, then the spatial arrangement of species determines which interactions dominate, thereby shaping their combined effect on community outcomes. We therefore asked whether our model could predict survival outcomes in communities with varying degrees of spatial mixing.

To address this question, we extended our reaction-diffusion model to two dimensions. We simulated chambers containing *P. aeruginosa* and *S. aureus* arranged in a checkerboard pattern with increasing patch sizes. For each spatial configuration, we computed the steady state concentrations of HQNO and rhamnolipids and predicted the total *S. aureus* survival probability across the chamber. We found that survival probabilities are low when species are in close proximity but show a sharp, nonlinear increase as patches grow larger, because the balance shifts from the short-ranged tolerance-reducing rhamnolipids to the long-ranged tolerance-enhancing HQNO (Fig. 4A).

**Fig. 4.**
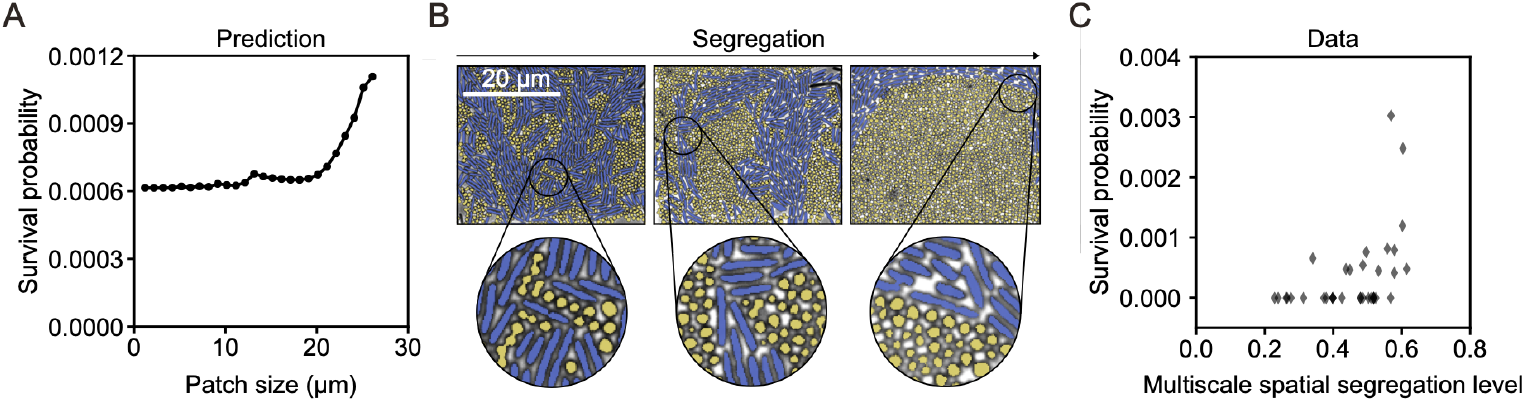
Spatial arrangement determines community-level tolerance. **(A)** Predicted *S. aureus* survival probability as a function of patch size from 2D reaction-diffusion model simulations, showing that survival increases non-linearly as patch size increases. **(B)** Representative images of growth chambers with increasing levels of spatial segregation. Images show segmented cells colored by species (*S. aureus:* yellow, *P. aeruginosa:* blue). Scale bar: 20 µm. **(C)** Experimentally measured *S. aureus* survival fraction as a function of spatial segregation (multiscale spatial segregation level, MSSL; see Methods). MSSL ranges from 0 for highly mixed to 1 for fully segregated arrangements. Survival increases significantly with increasing segregation (R^2^ = 0.31, p < 0.001, Spearman correlation). Each point represents one chamber (n = 40 chambers across 2 independent experiments).

To test the model predictions experimentally, we grew *S. aureus* and *P. aeruginosa* together in mixed communities (Fig. 4B). We measured the *S. aureus* survival fraction in each chamber and quantified the degree of spatial segregation between the two species using a previously published segregation index (36). Consistent with the model predictions, we found a positive correlation between segregation and survival fraction: chambers with greater separation between *P. aeruginosa* and *S. aureus* showed higher *S. aureus* survival (Fig. 4C). These results demonstrate that the interplay between HQNO and rhamnolipid concentration gradients determines *S. aureus* survival across both simple 1D geometries and more complex 2D spatial arrangements.

### Spatial arrangement modulates tolerance in colony biofilms

Having established that microscale spatial arrangement determines tolerance levels by balancing opposing interactions, we next asked whether similar mechanisms operate in three-dimensional colony biofilms. To control the spatial arrangement of colony biofilms, we adapted a microdroplet deposition platform (37) that spatially organized *S. aureus* and *P. aeruginosa* in predefined checkerboard patterns (Fig. 5A). The two species were arrayed on MHB agar pads at controlled center-to-center distances, creating checkerboards with small (111 μm), medium (222 μm), and large (555 μm) patch sizes (Fig. 5B). In addition, we created highly mixed biofilms by depositing the two species in overlapping areas. The communities were grown until the adjacent microcolonies merged into a single large biofilm. *S. aureus* and *P. aeruginosa* cell numbers were similar across all conditions (Fig. S16). Subsequently, the biofilms were dissociated, and tolerance levels were determined by plating.

**Fig. 5.**
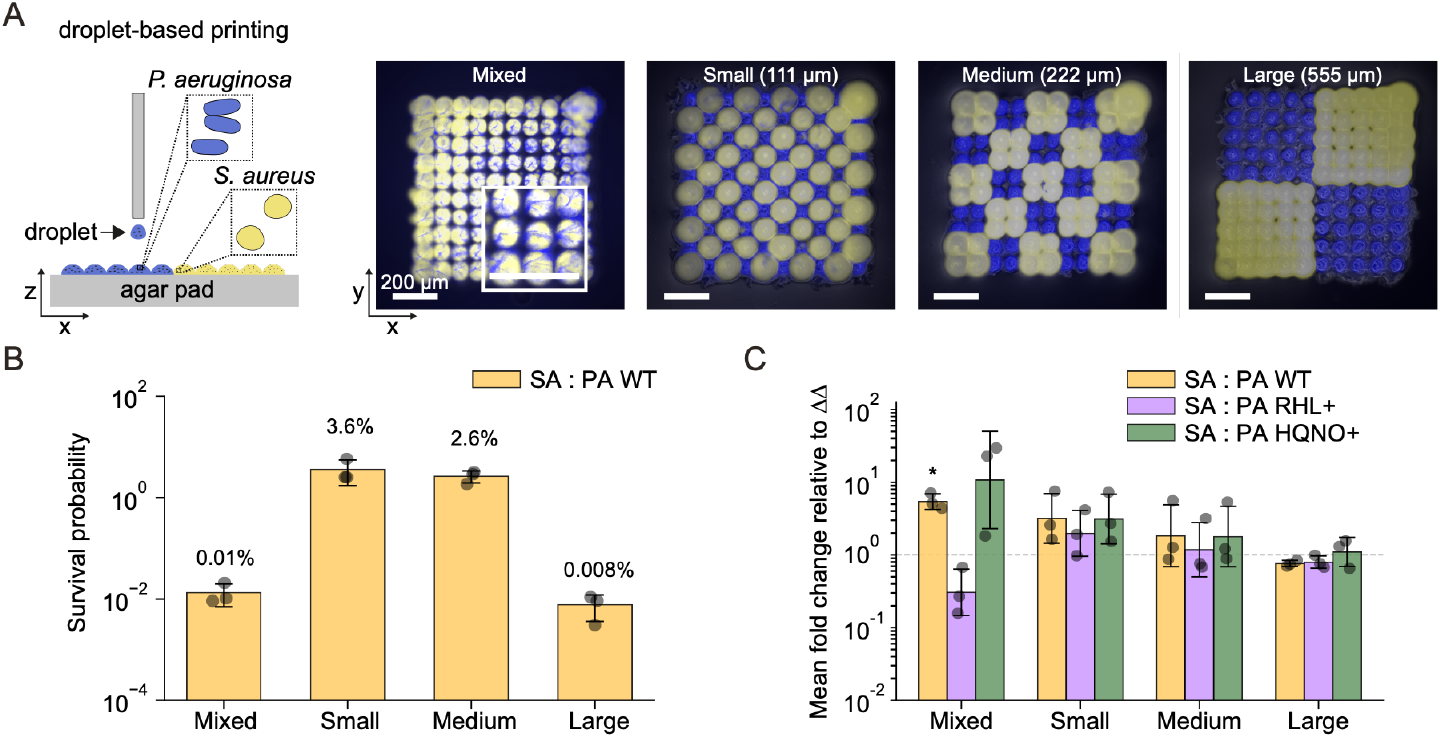
Spatial arrangement determines antibiotic tolerance in colony biofilms. **(A)** Biofilms with a checkerboard arrangement of *S. aureus* (yellow) and *P. aeruginosa* (blue) were printed on agar pads using droplet deposition. Representative fluorescence images of the four spatial arrangements with increasing patch size from left to right: mixed, small (111 μm), medium (222 μm), and large (555 μm). Scale bars: 200 μm. Inset shows higher magnification views. **(B)** *S. aureus* survival probability in co-culture with wild-type *P. aeruginosa* across the four spatial arrangements. Dots: individual replicates (n=3); bars: means; error bars: standard deviations. **(C)** Fold change in *S. aureus* survival in co-culture with *P. aeruginosa* strains producing different levels of HQNO and rhamnolipids, relative to co-culture with the *ΔpqsLΔrhlA* double mutant. Data shown for wild-type (yellow), HQNO-overproducing (HQNO+, green), and rhamnolipid-overproducing (RHL+, purple) *P. aeruginosa* across the four spatial arrangements. Dots: individual replicates (n=3); bars: geometric means; error bars: geometric standard deviations. *p < 0.05, one-sample t-test against baseline (log_10_ fold change = 0, dashed-line)

We observed that the biofilm’s spatial arrangement significantly influenced *S. aureus* tolerance levels (p = 0.04, Kruskal-Wallis test) (Fig. 5C). *S. aureus* survival was highest in biofilms with small and medium checkerboards (3.6% and 2.6%, respectively), but much lower in both mixed biofilms and large checkerboards (<0.01% for both).

To test whether HQNO and rhamnolipids also cause this pattern, we compared *S. aureus* survival in co-culture with *P. aeruginosa* strains producing different levels of these compounds. We used a double mutant lacking both compounds (*ΔpqsLΔrhlA*) as a baseline and compared it to the wild-type and HQNO- and rhamnolipid-overproducing strains. In the mixed biofilms, HQNO and rhamnolipids had the expected effects on *S. aureus* survival. Compared to the double mutant, *P. aeruginosa* wild-type and the HQNO-overproducing strain increased survival, while the rhamnolipid-overproducing strain reduced survival, though the effect was only significant for the wild-type (t-test, p < 0.05). In contrast, for the checkerboard patterns we found no significant differences, suggesting that HQNO and rhamnolipids do not strongly affect *S. aureus* tolerance in these biofilms. This likely reflects the short interaction ranges of these compounds relative to the checkerboard patch sizes.

These results show that HQNO and rhamnolipids affect antibiotic tolerance in *S. aureus* in mixed colony biofilms where the two species grow in close proximity. In checkerboard biofilms with greater spatial separation, arrangement strongly affects tolerance, but these patterns are not mediated by HQNO and rhamnolipids, suggesting additional *P. aeruginosa*-mediated interactions.

## Discussion

This study shows that community-level properties, such as antibiotic tolerance, emerge from the combined effect of multiple interactions. When interactions differ in their signs and ranges, they can generate complex spatial patterns that cannot be predicted from studying interactions in isolation. Because spatial arrangement modulates the relative contribution of these interactions, it is critical in determining community-level properties. Our framework thus shows that community-level properties are determined by three key factors: the relative strength and sign of interactions, their characteristic ranges, and the spatial arrangement of interacting species.

While individual interactions can structure tolerance spatially (29, 38), we show here that multiple opposing interactions can generate qualitatively different patterns. In *S. aureus* and *P. aeruginosa* communities, tolerance was highest at intermediate distances from *P. aeruginosa*. This non-monotonic pattern results from a dominant short-ranged negative interaction combined with a weaker long-ranged positive interaction (Fig. 1B). These differences can be explained by the underlying molecular parameters (18). The longer range of HQNO can be attributed to its higher diffusion coefficient and lower uptake rate (Fig. S14A). The dominance of rhamnolipids results from its higher effective concentration: although *S. aureus* is more sensitive to HQNO, this is compensated for by the higher production rate of rhamnolipids in *P. aeruginosa* (Fig. S14A). HQNO and rhamnolipids are typically considered antagonistic molecules with bacteriostatic and bactericidal activity against *S. aureus*, respectively. However, the concentrations in our chambers were subinhibitory, as *S. aureus* growth rates were similar at all distances from *P. aeruginosa* (Fig. S7).

We assumed that HQNO and rhamnolipids have additive effects on tolerance as they affect tobramycin tolerance through distinct mechanisms: HQNO leads to a reduction in the proton motive force (PMF) required for tobramycin uptake (24), while rhamnolipids mediate pore formation in the membrane, providing an alternative entry route (24). Although the additive model worked well for the current system, molecular compounds can have non-additive effects on cells (39). Such non-linearities would add a further layer of complexity and potentially allow for even more complex spatial patterns.

Our framework shows that spatial arrangement determines community-level properties by balancing multiple range-dependent interactions. Specifically, for tolerance, we predicted a peak at intermediate segregation: as segregation increases, dominance shifts from short-ranged rhamnolipids (low tolerance), through long-ranged HQNO (high tolerance), to neither interaction (low tolerance). Consistent with these predictions, tolerance in microfluidic chambers increased sharply with segregation (Fig. 4), and HQNO and rhamnolipids affected tolerance in mixed biofilms but not in larger checkerboard patterns (Fig. 5). However, technical limitations in bioprinting resolution (∼100 μm minimum spacing) prevented us from directly testing whether tolerance peaks at intermediate segregation in biofilms. These findings are also consistent with previous work showing that spatial structure enhances *S. aureus* tolerance in murine wound models and that this effect is eliminated by homogenization or HQNO deficiency (29).

The spatial scales at which HQNO and rhamnolipids affect tolerance could be relevant in clinical biofilms. In cystic fibrosis lungs and chronic wounds, aggregates of approximately 2–200 µm diameter form (40–42), and imaging studies reveal interspecies distances ranging from a few to several hundred micrometers (29, 40, 42–46). Our quantified interaction ranges of 32.8 µm for HQNO and 13.8 µm for rhamnolipids fall within this clinically relevant length scale, suggesting that the spatial tolerance patterns we observe could emerge *in vivo*. However, production levels of HQNO and rhamnolipids vary between *P. aeruginosa* isolates (24) and depend on environmental conditions, such as oxygen availability (47) or host immunity factors (33, 48). Similarly, *S. aureus* tolerance levels, and how they are modulated by external compounds, vary between conditions (49–51). Characterizing these interactions in vivo will be needed to test clinical relevance. Nonetheless, our findings add to a growing body of work highlighting the importance of considering microscale spatial arrangement (i.e., biogeography) in infections (52, 53).

We focused on two well-characterized compounds affecting antibiotic tolerance. However, *P. aeruginosa* secretes numerous other metabolites that affect *S. aureus* (54, 55), and two observations suggest additional compounds are relevant in this system. First, tolerance was reduced close to *P. aeruginosa* even without rhamnolipids (Fig. 2), suggesting an additional short-ranged tolerance-reducing interaction. Second, tolerance was higher in small and medium checkerboard biofilms than in both mixed and large checkerboards (Fig. 5). This effect was independent of HQNO and rhamnolipids, suggesting additional range-dependent interactions. Approaches such as (spatial) transcriptomics could identify these additional molecular players in future work.

A further limitation is that we focused solely on spatial dynamics and ignored temporal dynamics. Because tolerance is by definition a transient state, precise spatial patterns critically depend on treatment time. Moreover, while our microfluidic chambers reach steady state before antibiotic treatment, growing biofilms are more dynamic: production rates, sensitivities, and spatial arrangement could all vary with time. Integrating temporal dynamics represents an important future direction.

Our findings demonstrate that biofilm properties like antibiotic tolerance emerge from multiple interactions operating at different length scales and cannot be understood by studying interactions in isolation. Our framework provides a foundation for predicting how spatial arrangement modulates such interactions to shape community-level phenotypes.

## Materials and Methods

### Bacterial Strains and culture conditions

Bacterial strains used in this study are listed in Table S1. For *P. aeruginosa*, we used *PAO1 ΔpelA-ΔpslABC-ΔpilA-ΔfliC* as the background strain, which was previously optimized for use in microfluidics by removing matrix production and motility (56). We refer to this strain as *P. aeruginosa* wild-type throughout. For *S. aureus*, we used the methicillin-resistant strain BV18 USA300 (MRSA). All experiments were performed in Mueller-Hinton Broth (MHB). We chose this medium for consistency with previous studies in which tolerance-modulating interactions between *P. aeruginosa* and *S. aureus* were characterized (24, 25).

All batch cultures were grown at 37°C with shaking at 220 rpm. For *P. aeruginosa* strains carrying plasmids or chromosomal integrations, media were supplemented with appropriate antibiotics: chloramphenicol (30 µg/mL) for pSEVA331 transcriptional reporters, carbenicillin (100 µg/mL) for pQFT-Amp cumate-inducible expression reporters, and oxytetracycline (100 µg/mL) for mCTX::mNeonGreen chromosomal integrations. Strains carrying both pQFT-Amp plasmids and mCTX::mNeonGreen were maintained with carbenicillin (100 µg/mL) and oxytetracycline (100 µg/mL). For *S. aureus* strains carrying the pHC48 plasmid, chloramphenicol (10 µg/mL) was added to growth medium. *E. coli* strains were grown in Lysogeny Broth (LB) at 37°C with shaking at 220 rpm, with appropriate antibiotics added: chloramphenicol (30 µg/mL) for pSEVA331 plasmid, ampicillin (100 µg/mL) for pHC48 and pQFT plasmids, gentamicin (20 µg/mL) for pEX18Gm suicide vectors.

### Plasmids and oligonucleotides

Plasmids and oligonucleotides used in this study are listed in Tables S1 and S2, respectively. Plasmids were constructed using Gibson Assembly (57) and verified by Sanger sequencing. All plasmids were electroporated using a Bio Rad MicroPulser with preprogrammed settings: Ec1 for *E. coli* and *P. aeruginosa*, and Sta for *S. aureus*. All *P. aeruginosa* deletion mutants were generated by two-step allelic exchange using the suicide plasmid pEX18Gm (58, 59). For each target gene, approximately 800 bp homology regions flanking the deletion site were amplified from *P. aeruginosa* genomic DNA and assembled into pEX18Gm. Deletion mutants were confirmed by nanopore sequencing. Cumate-inducible expression vectors were constructed by first generating the pQFT-Amp backbone. AmpR was PCR-amplified from p2H12 and inserted into pQFT at the PvuII site. Target genes (*rhlAB* or *pqsL*) were then cloned into pQFT-Amp downstream of the synthetic PQJ promoter (60), enabling controlled overproduction of rhamnolipids or HQNO. The dual transcriptional reporter for monitoring *rhlA* and *pqsL* promoter activity was constructed by cloning the *rhlA* promoter region upstream of mcherry and the *pqsL* promoter region upstream of mNeonGreen into the pSEVA331 backbone (61), enabling simultaneous detection of rhamnolipid and HQNO biosynthesis gene expression. The *pflB* transcriptional reporter was constructed by cloning the *pflB* promoter region upstream of gfp in the pHC48 backbone (62), enabling detection of fermentative metabolism in *S. aureus* as a proxy for HQNO exposure. Fluorescent *P. aeruginosa* strains were generated using an integrative plasmid for chromosomal expression of mNeonGreen under control of a constitutive pX2 promoter, enabling visualization of *P. aeruginosa* in co-culture experiments. Primers used for plasmid construction and sequencing verification are listed in Table S2.

### Batch culture tolerance assay

To assess the effect of *P. aeruginosa* secreted factors on *S. aureus* antibiotic tolerance, we performed supernatant exposure assays. *P. aeruginosa* strains were grown in MHB for approximately 16 h. Cultures were centrifuged (4000 × g, 10 min) and supernatants were filter sterilized using 0.2 μm filters. *S. aureus* overnight cultures were diluted 1:100 in fresh MHB and grown to mid-exponential phase (OD600 ∼ 0.5). Cells were then treated with 30% (v/v) sterile *P. aeruginosa* supernatant and incubated for 30 min. This pre-exposure period was chosen to emulate conditions where bacteria encounter secreted metabolites prior to antibiotic treatment. Following pre-exposure, tobramycin was added at 58 μg/ml, similar to the Cmax achieved in humans at recommended dosing (63). This concentration is also within the range of sputum concentrations (17.2 to 327.3 μg/ml) reported for inhaled tobramycin therapy (64). An aliquot was collected before antibiotic addition for CFU enumeration (CFU_0_). After 8 h of treatment, an aliquot was removed and washed with PBS. Cells were serially diluted and plated on MHB agar to enumerate survivors. Survival fraction was calculated as CFU/CFU_0_.

### Microfluidic device fabrication

The microfluidic device consists of an array of adjacent chambers (50 × 50 × 0.80 μm^3^) flanked on opposite sides by flow channels measuring 22 μm in height and 100 μm in width. The chambers are connected by small gaps of 0.5 μm in width, 3 μm in length, and spaced by 2 μm. Nutrients enter each chamber from both sides, enabling a more uniform supply across the chamber length. The master mold was fabricated by direct laser writing into SU8, a negative photoresist that crosslinks where exposed to light, using a Heidelberg DWL66+ on two sequential spin coated layers (SU8-2001 and SU8-2025) (Pacheco, Ugolini et al., submitted). The microfluidic chips for the experiments were prepared by replica molding using Polydimethylsiloxane (PDMS, Sylgard 184, Dow Corning), mixed at a 1:5 ratio. The PDMS was cured at 80°C for 1 h, cut into chips of 3.5 cm × 3.5 cm, with 0.5 mm holes punched on both ends for loading cells and medium provision. For assembly, the PDMS chips and 22 x 44 mm diameter glass coverslips (VWR) were plasma treated for 1 minute using Harrick plasma cleaner PDC-32G, brought into contact, and then placed on a 100°C hot plate for bonding for 1 minute.

### Microfluidic device operation

Cultures were started from single colonies and grown overnight in 3 mL MHB. The following day, overnight cultures were diluted 1:100 in 5 mL growth medium and grown to mid-exponential phase (approximately 5 h for *S. aureus* and 2 h for *P. aeruginosa*). Cells were filtered through a 5 μm filter (Sartorius) to remove cell aggregates, then concentrated by centrifugation and resuspended in approximately 50 μL of growth medium. Prior to cell loading, microfluidic devices were pretreated by flushing with 1.0 mg/mL BSA in water to reduce cell adhesion to channel surfaces. After 1 minute, BSA was removed by flushing with growth medium.

*S. aureus* was loaded into chambers on one side of the porous wall and *P. aeruginosa* into chambers on the opposite side, by pipetting approximately 1 µL of concentrated cells into the respective flow channels. After loading, the flow channels were continuously flushed with growth medium at 0.2 mL/h using a syringe pump (Pump 11 Pico Plus Elite, Harvard Apparatus). A single 10 mL syringe was used to supply both flow channels by splitting the flow using a PEEK Y connector (Darwin Microfluidics) and connecting to the microfluidic device via PTFE tubing (0.56 mm ID × 1.07 mm OD, Fisher Scientific). Waste medium was collected via tubing connected to the outlet.

A metal connector positioned upstream of the Y junction enabled medium switching during experiments, by disconnecting tubing from the current syringe and connecting it to a new syringe containing the desired medium.

### Microfluidic tolerance assay

Cells were grown for approximately 16 h in MHB until chambers reached a dense monolayer. Subsequently, the medium was switched to MHB supplemented with 25 μM orange-red TAMRA-based fluorescent D-amino acid (RADA, Tocris) and 58 μg/mL tobramycin (Sigma Aldrich, 64× MIC) for 3 h. Afterwards, the medium was switched back to fresh MHB for 20 to 24 h to enable recovery.

### Microscopy and Image acquisition

Images were acquired with a Nikon Eclipse Ti2 Inverted Microscope equipped with a Hamamatsu ORCA-Flash4.0 V3 Digital CMOS camera (C13440-20CU) and a 100x NA1.45 Nikon Plan Apo Lambda Oil Ph3 DM objective (MRD31905). Fluorescence imaging was acquired with LED light at 470nm (for GFP) and 575nm (for RADA and PI) using a SPECTRA-X LED Illumination System as light source. Emitted light was filtered using a GFP/mCherry ET dual band Chroma filter (F58-019). Focus was maintained using the Nikon perfect focus system and the setup was controlled using the Nikon NIS-Elements software. The sample was maintained at 37°C using an Okolab incubator. Time-lapse images were acquired with intervals of 6 minutes. The data was saved locally in ND2 format and later converted to tiff for analysis.

### Automated image analysis

Time-lapse tiff stacks were analyzed using the MIDAP image analysis pipeline (version: 0.3.18, available at: https://github.com/Microbial-Systems-Ecology/midap). For each position, images were cropped to the growth chambers using MIDAP’s interactive ROI cutout tool. To avoid phase contrast artifacts near the flow channel, the ROI excluded the ∼8 µm closest to the channel edge. Cell segmentation was performed on the phase-contrast channel using a StarDist model which was previously trained on images acquired with the same microfluidic device (65). For each segmented cell, fluorescence intensity was quantified in the GFP and/or mCherry channels by computing the mean pixel intensity within the corresponding segmentation mask.

### Spatial analysis of regrowth events

Tolerant cells capable of regrowth dilute RADA through cell division and lose fluorescence, while non-regrowing susceptible cells retain the fluorescent label. As regrowing cells divide, their offspring form clusters of actively growing cells. To count only unique independent regrowth events, we implemented an automated analysis pipeline. We first classified cells as growing or non-growing based on their RADA fluorescent intensities using a dynamic threshold. To distinguish fluorescent (growing) from non-fluorescent (non-regrowing) cells, we applied an automated threshold based on kernel density estimation (KDE). We computed a KDE of the normalized fluorescence intensity distribution, then used peak detection (scipy.signal.find_peaks) to identify local minima (valleys) and maxima (peaks). The threshold was defined as the midpoint between the first valley and the highest peak to its left, effectively separating the two populations in the bimodal distribution. Cells with fluorescence above the threshold were classified as growing; those at or below were classified as non-regrowing. The resulting binary images were processed to merge adjoining regrowing cells into continuous clusters using a binary expansion operation with a radius of 7-pixels. We then tracked these clusters over time by comparing consecutive frames using overlap-based tracking. Unique regrowth events were defined as the appearance of new clusters. For the *ΔpqsL* and *ΔpqsLΔrhlA* mutant conditions, most chambers showed no regrowth, and in the remaining chambers, only individual cells regrew. These data were therefore analyzed manually to avoid the computational expense of the automated pipeline.

### Fluorescent Reporter Calibration

To quantify HQNO and rhamnolipid (RHL) concentrations in microfluidic experiments, we used fluorescent reporters and constructed calibration curves relating fluorescent intensities to compound concentrations. We first measured dose-response curves by exposing *S. aureus* reporter strains to a range of purified HQNO and RHL concentrations using flow cytometry. We then repeated measurements at selected concentrations in the microfluidic system to calibrate fluorescence intensities between the two measurement modalities.

#### Reporters

For HQNO, we used a GFP transcriptional reporter for *pflB* expression, a *pflB* promoter-gfp transcriptional fusion that uses *pflB* activity as a proxy inhibited respiration induced by HQNO. For RHL, we followed a previously published protocol that uses uptake of propidium iodide (PI) as a proxy for membrane damage.

#### Flow cytometry dose-response curves

To measure the HQNO dose-response, *S. aureus* exponential phase cultures carrying *pflB*-gfp were exposed to purified HQNO (Focus Biomolecules) for 8 h in culture tubes. GFP expression was measured with 488 nm excitation and 502–525 nm emission.

For RHL, wild-type *S. aureus* exponential phase cultures were exposed to purified RHL (Sigma Aldrich) for 8 h, following addition of 100 µg/mL PI for 30 minutes. PI fluorescence was measured with 561 nm excitation and 595–654 nm emission.

At least 10,000 events per sample were recorded by flow cytometry (Fortessa, BD Biosciences). Data were analyzed using FlowJo software (version 10.10.0). Per cell fluorescent intensities were background-corrected using the median cell intensity for the 0 concentration controls, and the median was taken over all cells per replicate. Each condition was measured in triplicate.

#### Microfluidic calibration measurements

For HQNO calibration, *S. aureus pflB*-gfp was grown overnight in co-culture with *P. aeruginosa* Δ*pqsL*. Defined concentrations of HQNO were then added to both flow channels. Fluorescent intensities were measured after 8 h using microscopy settings described above.

For RHL calibration, wild-type *S. aureus* was grown overnight in co-culture with *P. aeruginosa* Δ*rhlA*. Defined concentrations of RHL were then added to both flow channels for 8 h. PI was added at 100 µg/mL for 30 minutes, and extracellular PI was subsequently flushed out to remove unbound dye. Fluorescence of intracellular PI was measured using microscopy setting described above.

All microscopy settings were held constant between experiments to enable quantitative comparisons. Images were segmented and per-cell fluorescent intensities were extracted as described above. Per-cell fluorescent intensities were background-corrected using the modal intensity in the 0 concentration control experiments. For each chamber, we used the median intensity across all cells as a representative fluorescent intensity, and each condition was measured in 10 chambers.

#### Cross-platform calibration

To convert flow cytometry fluorescence intensities to equivalent microscopy intensities, we performed a linear regression to calibrate the relationship between the two measurement modalities using samples at matched inducer concentrations. For HQNO, this calibration was performed using concentrations of 40, 80, 160, 320, and 2,500 ng/mL. For RHL, we used 0, 25, 50, 75, and 100 μg/mL; the measurement at 12.5 μg/mL was excluded as it was a clear outlier. Linear regressions were performed using Orthogonal Distance Regression to account for measurement uncertainty in both flow cytometry and microscopy data, using the SciPy ODR function. The regression analysis showed that microfluidic and microscopy intensities have linear relationships, with an R^2^ of 0.86 for HQNO and 0.98 for RHL (Fig. S12A,C).

#### Dose-response fitting

For RHL, fluorescent intensities measured using flow cytometry were converted into microscopy units using the linear regression, and dose-response curves were then fitted to the converted data. For HQNO, we also used converted flow cytometry measurements for HQNO concentrations of 40 ng/mL and above. However, for lower concentrations, fluorescent intensities were below the limit of detection of flow cytometry. For this range, we thus used the microscopy measurements instead.

Both HQNO and RHL reporters showed sigmoidal dose-response curves that could be approximated using a Hill function (Fig. S12B,D). Specifically, we fitted a three-parameter Hill function of the form 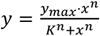 to the full concentration range, where *y* is fluorescent intensity, *x* is concentration, *y*_*max*_ is maximum intensity, *K* is the half-maximal concentration, and *n* is the Hill coefficient.

Although Hill functions fit the overall response well, they failed to accurately capture the dose-response relationship at the low intensities and concentrations that were relevant in our setup. To increase the accuracy of concentration estimates, we therefore re-fitted the low-concentration regime using a spline fit. We used a monotonic PCHIP (Piecewise Cubic Hermite Interpolating Polynomial) spline interpolator on log_10_-transformed concentration values using the SciPy PchipInterpolator function. For HQNO, we fitted up to a concentration of 160 ng/mL; for RHL, up to 100 μg/mL.

### Concentration Gradient Inference

To infer HQNO and RHL concentrations during co-culture with wild-type *P. aeruginosa*, we converted fluorescence gradients measured with *S. aureus* reporter strains into concentrations using the dose-response relationships.

#### Experiments

For HQNO measurements, *S. aureus pflB*-gfp was grown overnight in co-culture with *P. aeruginosa* wild-type, and GFP expression was measured after 16 h. For RHL measurements, *S. aureus* wild-type was grown overnight in co-culture with *P. aeruginosa* wild-type for 16 h. PI (100 µg/mL) was then added for 30 minutes, subsequently flushed out, and PI fluorescence was measured. All measurements used the same microscopy settings as the calibration experiments.

#### Image analysis

Images were segmented and per-cell fluorescent intensities were extracted as described above. Subsequently, per-cell fluorescent intensities were background-corrected using the modal intensity of the reporter strains in the 0-concentration calibration experiment. Cells were then spatially binned in fixed 1.5 μm bins as a function of distance from the *P. aeruginosa*-*S. aureus* interface, based on their centroid positions. For each bin, the median value over all cells was taken. Fluorescent intensities for each bin were then converted to concentrations using the inverse of the dose-response curves.

### Reaction-Diffusion Model

To quantify the strength and spatial range of interspecies interactions and predict survival outcomes in spatially mixed communities, we developed a mathematical model. Because compound concentrations equilibrate rapidly relative to cell growth, we implemented a steady state reaction-diffusion model to describe HQNO and RHL concentration gradients. We assumed constant production of HQNO and RHL by *P. aeruginosa* and constant uptake by *S. aureus*.

Specifically, we solved the following equation:

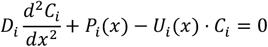

where *D*_*i*_ is the diffusion constant and *C*_*i*_ is the compound concentration; the subscript *i* denotes either HQNO or RHL. *P*_*i*_(*x*) is the spatially dependent production term, which is *P*_*i*_(*x*) = *p*_*i*_ in regions occupied by *P. aeruginosa* and 0 elsewhere. *U*_*i*_(*x*) is the spatially dependent uptake term, which is *U*_*i*_(*x*) = *u*_*i*_ in regions occupied by *S. aureus* and 0 elsewhere. For notational simplicity, we drop the *i* subscript in further equations.

We solved this equation for three different geometries.

#### 1D Receiver-only chamber

We first solved the steady state reaction-diffusion equation within the *S. aureus* chamber only. Given the symmetry across chamber width, we can solve a 1D version of the problem. In this region, the equation simplifies to:

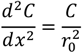

where 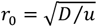 is the characteristic length scale of the reaction-diffusion process. We refer to this length scale as the *interaction range*.

We applied an absorbing boundary at the chamber end (*C*(*L*) = 0, where *L* is the length of the chamber), reflecting the large volume and high flow rate in the flow channel, and a fixed concentration boundary at the porous wall separating *P. aeruginosa* and *S. aureus* (*C*(0) = *c*_0_).

We solved this equation analytically to:

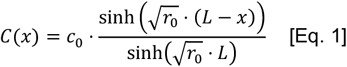

This equation was fitted to the experimentally inferred concentration gradients to infer *c*_0_ and *r*_0_ (see below).

#### 1D coupled chambers

Next, we solved the steady state reaction-diffusion equation for the full coupled chamber geometry. Given the symmetry across chamber width, we can solve a 1D version of the problem. The equation was solved separately in the following three domains:

Domain 1: *P. aeruginosa* producer chamber, (0, *L*):

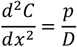

Domain 2: porous wall, (*L, L* + *d*), where *d* is the thickness of the porous wall:

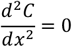

Domain 3: *S. aureus* receiver chamber, (*L* + *d*, 2*L* + *d*):

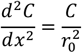

We imposed absorbing boundary conditions at the outer ends where the chambers exit into the flow channels: *C*(0) = 0 and *C*(*L* + 2*d*) = 0. At the interfaces between the domains, we enforced continuity of concentration (*C*_*left*._ = *C*_*right*_) and continuity of total-flux:

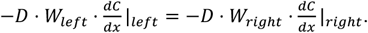

Here, *W*_*left*._ and *W*_*right*_ represent the cross-sectional widths to the left and right of the interface, respectively. These parameters account for the different diffusional cross-sections in the growth chambers (width *W*) versus the porous wall (combined gap width *W*_*gaps*_). The ratio *α* = *W*_*gaps*_/*W* reflects the effective porosity of the porous wall and quantifies the geometric reduction in inter-chamber diffusive transport.

We obtained analytical solutions using the Python SymPy library (66) (version 1.14.0) by solving the differential equations independently in the three domains and using the boundary conditions to solve for the integration constants.

To infer the production rate *p*, we used the following closed-form expression relating production rate *p* to the compound concentration at the receiver-side interface of the gap (*c*_0_):

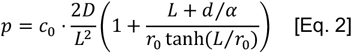

We solved the problem a second time with *p* = 0 to find the concentration *c*_*in*_ = *C*(0) that should be added to the flow channel at *x* = 0 to achieve the same concentration *c*_0_ at the interface for the non-producing *P. aeruginosa* double mutant:

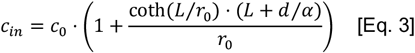

#### 2D coupled chambers

To study the effect of interactions in mixed communities, we solved the full 2D model numerically using an explicit finite-difference scheme with a square grid (Δx = Δy = 1 μm). The discretized domain captured the full geometry of the coupled chambers, including the pillars forming the porous wall. We imposed absorbing boundary conditions at the outer chamber ends (where chambers exit into flow channels) and reflecting boundary conditions at all other boundaries, including pillar surfaces.

We computed the Laplacian operator using a 5-point stencil and performed time-stepping with an adaptive time step satisfying the CFL stability criterion. Simulations were run until convergence (maximum concentration change < 10^−6^ per time step). The simulations were implemented in Python using NumPy and were optimized using Numba JIT compilation.

### Parameter Inference

Model parameters (Table S3) were obtained from literature or directly estimated from experiments where possible. All other parameters were inferred by fitting the model to the data.

#### Diffusion coefficients

The diffusion coefficient of RHL, *D*_*RHL*_ = 27 μm^2^/s, was obtained from literature (67). For HQNO, no literature estimates could be found, and we instead used the theoretical diffusion parameter calculated using the Stokes-Einstein-Sutherland equation. Based on HQNO’s molecular weight of 259.34 g/mol and assuming a spherical molecular shape, we estimated a molecular radius of 0.5 nm, yielding *D*_*HQNO*_ = 40 μm^2^/s.

To account for hindered diffusion in densely populated microfluidic chambers, we applied a porous-media correction: 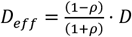, where *ρ* denotes the cell volume fraction (68). We estimated the 3D cell density from segmented phase-contrast images. We first estimated the 2D area fraction (*ρ*_2*D*_) by taking the fraction of pixels occupied by cells relative to the total number of pixels per chamber. This was converted to a 3D volume fraction (*ρ*) using the cell radius (*r*_*cell*_) and chamber height (*h*), according to 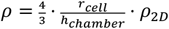. Averaging across ten chambers gave *ρ*_2*D*_ = 0.42 and *r*_*cell*_ = 0.325 μm; using the measured chamber height of *h* = 0.8 μm, this corresponds to *ρ* = 0.463.

#### Interaction range and maximum concentration

The interaction range *r*_0_ and maximum concentration *c*_0_ were inferred simultaneously by fitting the theoretical concentration profiles (Eq. 1) to the experimentally inferred concentration profiles. Nonlinear least-squares fitting was done using the SciPy curve_fit function with constraints *c*_0_ ≥ 0, *r*_0_ > 0, and *L* ≥ *L*_*chamber*_ + 1.5 μm. *L* was treated as a free parameter, as the microscopy images were cropped to exclude the chamber-flow-channel interface to reduce optical artifacts.

RHL was fitted over the domain 1.5 ≤ *x* ≤ 45 μm. The first bin next to the porous wall and last bins close to the flow channel were excluded to remove edge effects. HQNO was fitted over the domain 7.5 ≤ *x* ≤ 45 μm. Additional bins next to the porous wall needed to be excluded, as *pflB* reporter levels were lower here than in the center of the chambers, as additional factors reduced expression levels close to *P. aeruginosa*. We therefore constrained the fit to the region where *pflB* decreased monotonically with distance. The precise cut-offs were chosen based on maximizing the *R*^2^ values of the fits.

#### Uptake and production rates

Uptake rates were directly calculated from the interaction range *r*_0_ using the relation 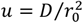. Production rates were calculated analytically using Eq. 2 from the estimated values of the maximum concentration at the *S. aureus* side of the porous wall (*c*_0_) and the interaction range (*r*_0_). To account for correlations in parameter estimation, *c*_0_ and *r*_0_ estimated from the same chamber were treated as paired values, and a single production rate was estimated for each chamber individually. The geometric parameters *L* = 50 μm, *d* = 3 μm, and *α* = 0.25 were obtained from the design files of the microfluidic device.

### Survival Model

We developed a quantitative model relating *S. aureus* survival to HQNO and RHL concentrations using batch culture survival assays.

#### Experimental data

We grew *S. aureus* to mid-exponential phase in 1 ml of MHB medium within 24-well plates and subsequently added HQNO and RHL at defined concentrations. After a 30-minute incubation period, an aliquot was collected for CFU counting and tobramycin at 58 μg/mL was added. After 3 h of treatment, survival was assayed using CFU counting.

#### Model fitting

We fitted a logistic regression model to survival data with the form:

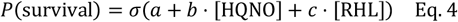

where 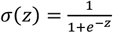 is the logistic function, *b* and *c* are coefficients quantifying the effect of each molecule on survival, and *a* is the intercept term. We validated that this linear model outperformed models with quadratic and/or interaction terms.

To optimize performance, the model was fitted over an HQNO range of 0–60 ng/mL, which corresponds to the range of concentrations inferred in the microfluidic chambers. Data points below the limit of detection were excluded from fitting. As growth during early antibiotic treatment could lead to survival >1, all survival values were normalized by dividing by the maximum observed survival. Subsequently, the model was fitted using least-squares regression on logit-transformed survival values using the SciPy curve_fit function.

#### Survival prediction

We combined predicted concentration profiles with the logistic survival model to estimate spatially resolved survival probabilities. Model uncertainty was evaluated by exhaustively sampling all combinations of fitted HQNO and RHL concentration profiles. Specifically, we sampled all combinations of (*c*_0,HQNO,*i*_, *r*_0,HQNO,*i*_) and (*c*_0,RHL,*j*_, *r*_0,RHL,*j*_), where *i* and *j* refer to chamber index, and calculated HQNO and RHL concentration profiles using Eq. 1. We then applied Eq. 4 to estimate the survival probability from these concentration profiles. This procedure accounts for possible correlations between *c*_0_ and *r*_0_ parameters inferred from the same chamber.

### Complementation Assay

To validate our model, we aimed to complement the non-producing Δ*pqsL*Δ*rhlA P. aeruginosa* double mutant by adding purified HQNO and RHL exogenously to the flow channel on the *P. aeruginosa* side. These concentrations were estimated using Eq. 3, taking the median estimated value. *S. aureus* was grown in co-culture with the Δ*pqsL*Δ*rhlA P. aeruginosa* double mutant following the protocols detailed above. The media on the *P. aeruginosa* side consisted of MHB supplemented with 207 ng/mL HQNO and 200 μg/mL RHL, while the media on the *S. aureus* side consisted of MHB. After overnight growth, tolerance assays were performed as described above.

### Mixed Chamber Survival

#### Model predictions

We used the full 2D model to predict how segregation in mixed communities affects survival. Matching the experimental setup, one chamber was left empty, while the other connected chamber was filled with a mixed community of *S. aureus* and *P. aeruginosa* using a checkerboard pattern. The size of the checkerboard patches was systematically varied, from 1 μm (1 cell per patch) to 25 μm (4 patches in total, with 625 cells each). We then ran the 2D model to find steady state concentration profiles of HQNO and RHL and used the logistic model (Eq. 4) to predict survival. We then calculated the average survival probability of *S. aureus* by averaging over all cells in the chamber.

#### Experimental setup

Microfluidic tolerance assays were performed as described above, with one major change: *S. aureus* and *P. aeruginosa* were loaded from one side, leaving the other side uncolonized. While chambers show high degrees of mixing directly after loading, the species naturally segregate as the community grows. To vary the degree of segregation, we thus combined two experiments. For high mixing, *S. aureus* and *P. aeruginosa* were loaded simultaneously and incubated for 2 h before treatment. For high segregation, *S. aureus* was loaded 4 h before treatment and *P. aeruginosa* was added 2 h before treatment, allowing additional time for spatial separation. Both conditions thus provided 2 h of *S. aureus* contact with *P. aeruginosa* prior to antibiotic treatment. We then selected 15-20 chambers for each condition, while aiming to maximize the diversity of spatial arrangements. For each chamber, we quantified survival at the single-cell level as described above and calculated the average survival frequency per chamber.

#### Segregation score calculation

To quantify spatial segregation between producers and consumers, we used the multiscale segregation analysis developed by Dogsa & Mandic-Mulec (2023). This method calculates the degree of segregation in a local window of varying size. For each window size, the segregation index for *S. aureus* is given by:

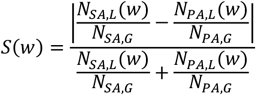

where *N*_*SA,L*_(*w*) and *N*_*PA,L*_(*w*) are counts of *S. aureus* and *P. aeruginosa* cell types within distance *w* (window size) from the focal *S. aureus* cell, and *N*_*SA,G*_ and *N*_*PA,G*_ are the corresponding global counts of each type. For the simulations, we used Chebyshev neighborhoods with a radius of 1 to 25 pixels (corresponding to 1 to 25 μm). For the experiments, we used Euclidean center-to-center distances, with a neighborhood radius of 2 to 25 μm.

To capture segregation across spatial scales, we followed Dogsa & Mandic-Mulec (2023) and computed the MultiScale Segregation Level (MSSL) score by calculating the area under the segregation-window size curve, normalized by the window size range:

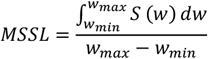

where the integral was computed using the trapezoidal rule using the SciPy trapezoid function, after excluding windows for which *S*(*w*) was undefined.

### Colony Biofilms

Spatially defined dual-species biofilms were generated using a computer-controlled pressure pump system that deposited liquid samples from microcentrifuge tubes through a hydrophobic capillary onto a single 3% agarose pads cast in a glass-bottom petri dishes. Three checkerboard patterns of *S. aureus* pHC48.dsRed and *P. aeruginosa* mNeonGreen were printed with interspecies segregation distances ranging from 110 µm to 500 µm. In addition, a mixed biofilm pattern was created by depositing the two strains in the same location.

The agar pads were incubated at 37°C for approximately 8 h to permit colony formation. Checkerboard patterns were then excised from the agarose and vortexed in 1 mL MHB in 2 mL microcentrifuge tubes. A 10 µL aliquot was collected for serial dilution and CFU enumeration prior to antibiotic treatment. Tobramycin was added to a final concentration of 58 µg/mL, and tubes were incubated at 37°C with shaking at 600 rpm for 1 h. To minimize *S. aureus* cell loss during washing steps required for antibiotic removal, cultures were supplemented with approximately 10^9^ CFU of the diaminopimelic acid (DAP)-dependent *E. coli* strain JKe201 (69). This carrier biomass enabled formation of a stable cell pellet during centrifugation and PBS washing. Resuspended pellets were plated on Baird-Parker agar (selective for *S. aureus*) and LB agar containing 100 µg/mL oxytetracycline (selective for *P. aeruginosa*), both lacking DAP supplementation, thereby permitting enumeration of *S. aureus* and *P. aeruginosa* while preventing growth of the helper *E. coli* strain.

### Statistical Analysis

All analyses were performed in Python 3.9.10 using NumPy 1.24.4 (70), SciPy 1.13.1 (71), and Pandas 1.5.3 (72), while Matplotlib 3.5.1 (73) and Seaborn 0.13.2 (74) were used for plotting. Statistical methods are detailed in figure captions.

For statistical purposes, we considered individual growth chambers as independent biological replicates. This assumption is justified by the comparatively large volume and high flow rate in the flow channels, which ensure that compounds produced in one chamber have negligible effects on downstream chambers. Consequently, interactions and spatial pattern formation develop independently in each growth chamber. All experiments were replicated across at least 2 independent microfluidic devices. Data from chambers across different devices were pooled after confirming by visual inspection that there were no systematic differences between devices.

## Data and Code Availability

Figure source data files and Jupyter notebooks containing the analysis code for the experimental data are available on GitHub: https://github.com/simonvanvliet/Spatial-Tolerance-Figure-Data-and-Code.

The model code, figure source data files, and Jupyter notebooks with analysis code for the model-related figures are available on GitHub: https://github.com/simonvanvliet/SpatialToleranceModel.

Raw imaging data and the output from segmentation and tracking analysis will be made available on BioImageArchive during revision.

## Acknowledgements

GitHub Copilot was used as a coding assistant, and Claude 4 Sonnet (Anthropic) was used for coding advice and language editing. All AI-generated content was critically reviewed, verified, and edited by the authors. Calculations were performed at sciCORE (http://scicore.unibas.ch/) scientific computing center at University of Basel.

We thank Knut Drescher and members of the Drescher and van Vliet labs for valuable feedback and suggestions. We also thank Alyssa Henderson for valuable feedback on the manuscript, Ashley Alexander and Joanna Goldberg for providing the pHC48 plasmid, Julian Bär and Annelies Zinkernagel for the trained StarDist model, and Maya Bruderer for assistance in cloning *P. aeruginosa* mutants.

## Author Contributions

Conceptualization: GB, SvV

Methodology: GB, BR, LF, GSU, SvV

Formal analysis: GB, BR, SvV

Investigation: GB, LF

Resources: JK, AK, ATA, GSU, RS, UJ, DB, PSD

Writing – Original Draft: GB, BR

Writing – Review & Editing: GB, OTS, SvV

Visualization: GB

Supervision: RS, UJ, DB, PSD, OTS, SvV

Project administration: SvV

Funding acquisition: SvV

