## Supplementary material for "Opposing Range-Dependent Interactions Create Complex Spatial Patterns of Antibiotic Tolerance in Multispecies Biofilms": SI Figures and Tables

### Supplementary Information

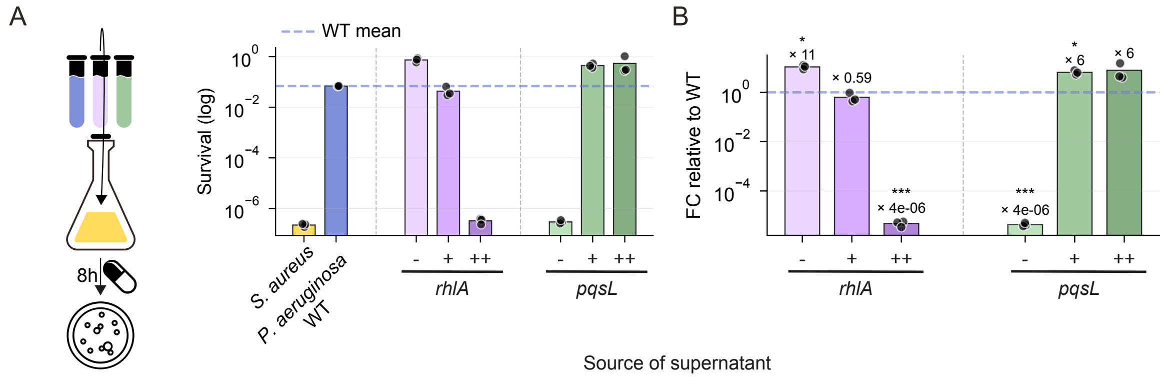


**Figure S1. Rhamnolipids and HQNO modulate tobramycin tolerance in *S. aureus*.** *S. aureus* was grown to mid-exponential phase in MHB medium and pre-treated with sterile supernatants (30% v/v) from *P. aeruginosa* strains PAO1 wild-type or its isogenic mutants (-), complemented (+) or overexpressing strains (++) for 8 h. Subsequently, an aliquot was plated to determine baseline CFU count ($CFU_0$), and the remaining cells were treated with 58 µg/ml tobramycin for 8 h. **(A)** Survival fraction ($CFU/CFU_0$) at 8h post-antibiotic treatment, shown on a $log_{10}$ scale. **(B)** Fold change in survival relative to *P. aeruginosa* wild-type (WT) (dashed line). Bars represent means; individual points show biological replicates. Statistical analysis: one-sample t-test on $log_2$-transformed fold changes with Bonferroni correction. *p < 0.05, **p < 0.01, ***p < 0.001; n = 3.

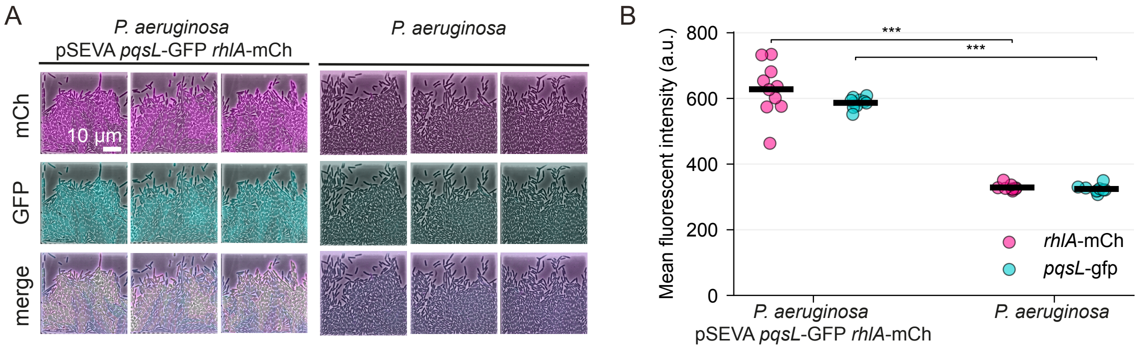


**Figure S2. Transcriptional activity of HQNO and rhamnolipid biosynthesis genes. (A)** Representative fluorescence microscopy images of *P. aeruginosa* + pSEVA *pqsL*-GFP *rhlA*-mCherry, carrying a plasmid-based dual transcriptional reporter for HQNO production (*pqsL*) and RHL production (*rhlA*), and wild-type *P. aeruginosa* (autofluorescence control) grown in microfluidic chambers. Scale bar: 10 µm. **(B)** Mean fluorescence intensity of mCherry (magenta) and GFP (cyan) in reporter (n = 10) versus control (n = 9) strains. Each point represents the mean intensity per field of view; horizontal bars indicate group means. Statistical comparison by two-sample t-test; *p < 0.05, **p < 0.01, ***p < 0.001.

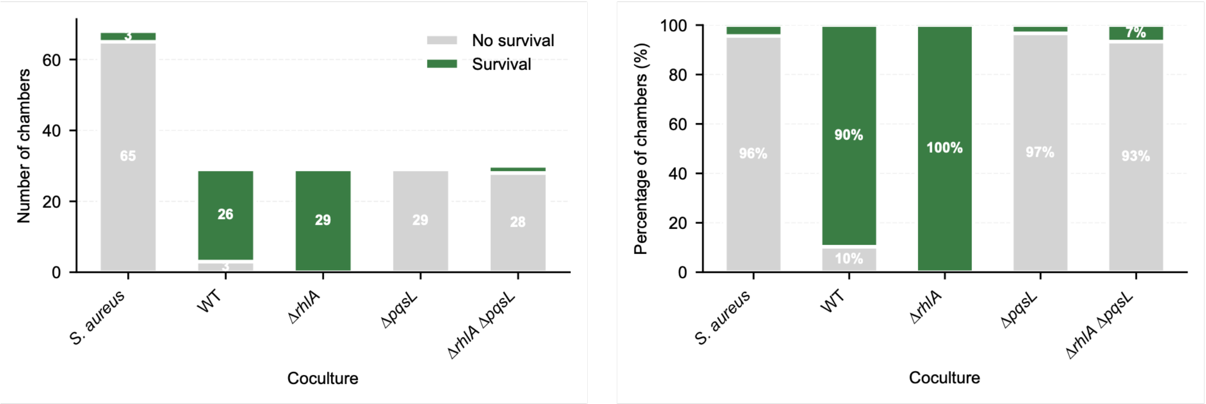


**Figure S3. *S. aureus* survival across cocultures. (A)** Total number of chambers scored for *S. aureus* survival (green) or no survival (grey) across different cocultures: *S. aureus* – *S. aureus* coculture, and *S. aureus* in coculture with *P. aeruginosa* WT or mutants (*ΔrhlA*, *ΔpqsL*, and *ΔrhlA ΔpqsL*). **(B)** Percentage of chambers showing *S. aureus* survival for each condition. Stacked bars represent the proportion of chambers with survival (green) versus no survival (grey) outcomes. Data represent 68 chambers (*S. aureus* – *S. aureus* coculture), 29 chambers (WT and *ΔrhlA* cocultures), and 30 chambers (*ΔpqsL* and *ΔrhlA ΔpqsL* cocultures) analyzed across independent microfluidic devices.

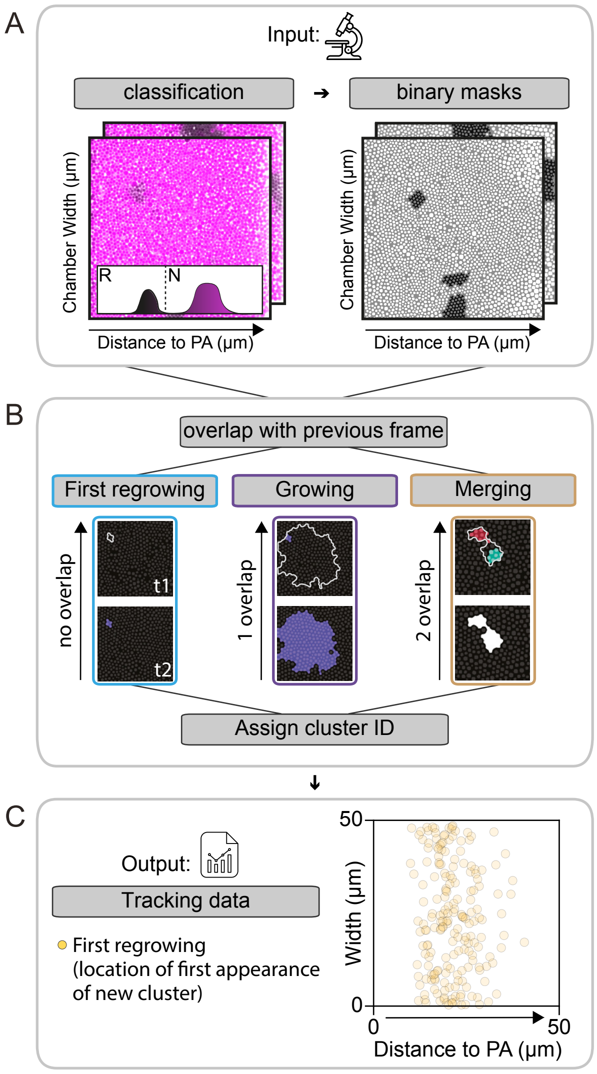


**Figure S4. Image analysis pipeline for tracking *S. aureus* regrowth in microfluidic chambers. (A)** Microscopy images of *S. aureus* chambers post-tobramycin removal, overlaying phase contrast (grey scale) with RADA fluorescence (magenta). Cells are classified as regrowing (R; lose fluorescence by diluting RADA through growth) or non-regrowing (N, remain stained) by thresholding on per-cell fluorescence intensities, resulting in binary masks. **(B)** The binary masks from sequential timepoints are compared to detect overlap. Clusters are classified as: "First regrowing" (new cluster with no overlap to previous frame), "Growing" (single cluster overlap, indicating continued growth), or "Merging" (two cluster overlaps, indicating clusters merged). Each cluster is assigned a unique identifier. **(C)** Output data showing the spatial coordinates of first regrowing clusters, plotted as distance from *P. aeruginosa* (PA) versus chamber width.

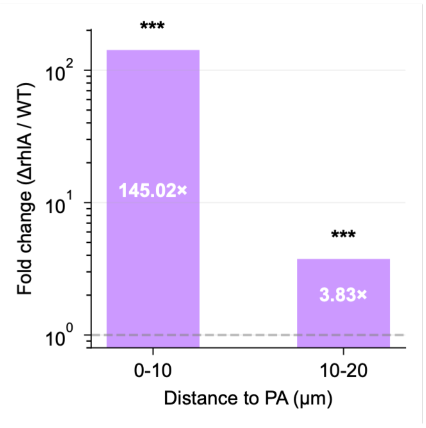


**Figure S5. Distance-dependent effect of rhamnolipids on regrowth.** Fold change in regrowth fraction (*ΔrhlA* / WT) at 0–10 μm and 10–20 μm distance from *P. aeruginosa*. Within each distance bin, WT and Δ*rhlA* regrowth fractions were compared using Mann-Whitney U test (*p < 0.05, **p < 0.01, ***p < 0.001). Bars show the fold change calculated as mean regrowth fraction in *ΔrhlA* divided by mean regrowth fraction in WT. Dashed line indicates fold change = 1 (no difference).

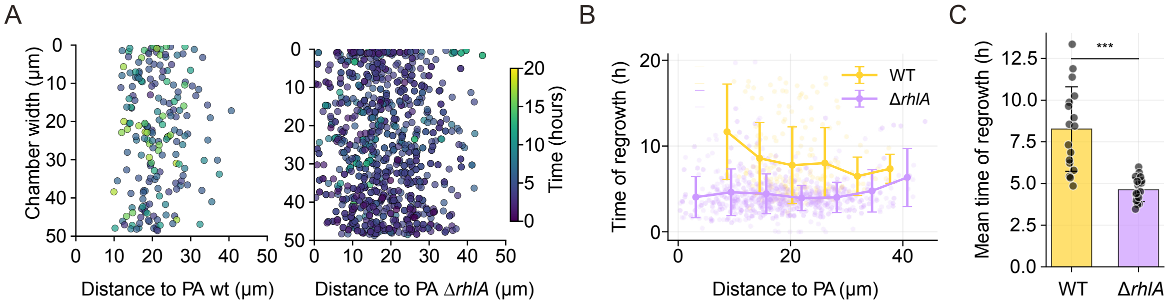


**Figure S6. *S. aureus* regrowth after antibiotic treatment occurs faster in absence of rhamnolipids. (A)** Spatial distribution of first regrowing *S. aureus* clusters in microfluidic chambers with wild-type *P. aeruginosa* (left) or Δ*rhlA* mutant lacking rhamnolipid production (right). Each point represents one regrowing cluster; color indicates time to regrowth after tobramycin treatment. **(B)** Regrowth time as a function of distance from *P. aeruginosa*. Lines connect binned means (error bars: SD); individual clusters shown as transparent points. **(C)** Bars show mean regrowth time averaged across chambers (error bars: SD). Each point represents one microfluidic chamber. *p < 0.05, **p < 0.01, ***p < 0.001, two sample t-test on chamber-level means.

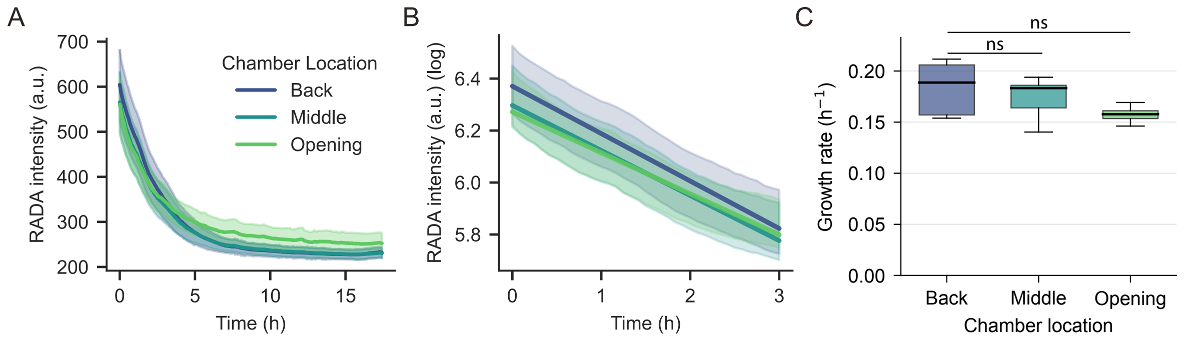


**Figure S7. *S. aureus* growth rate is uniform across microfluidic chamber**. *S. aureus* was pre-labelled with the fluorescent D-amino acid RADA, then switched to RADA-free medium. As cells grow, newly synthesized unlabeled peptidoglycan dilutes the RADA signal; the rate of fluorescence decay was used to infer cell growth rate. **(A)** Per-cell RADA fluorescence intensity over time for three spatial bins measured from the porous wall: Back (0-15 µm), Middle (15-30 µm), and Opening (30-45 µm). Lines show mean; shaded regions indicate SD. **(B)** Log-transformed RADA intensity for the same spatial bins as in (A) during the first 3 hours; linear decay indicates exponential growth. **(C)** Growth rates calculated from fluorescence decay slopes. Statistical analysis: One-way ANOVA; ns = not significant.

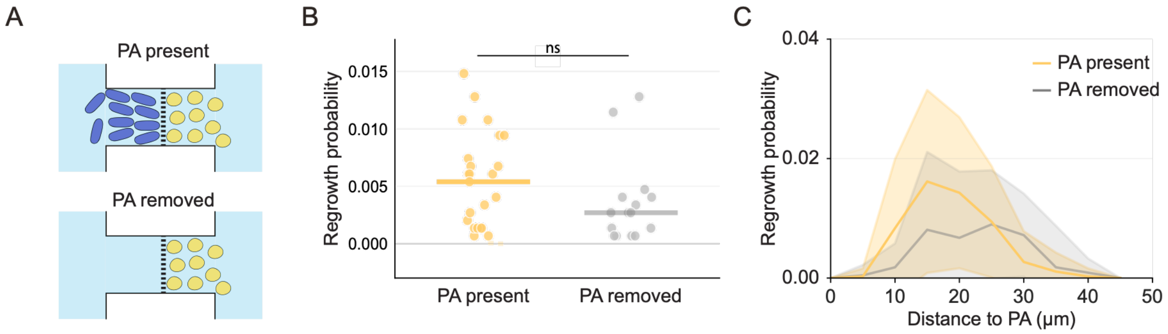


**Figure S8. Removal of *P. aeruginosa* does not alter *S. aureus* regrowth probability. (A)** *P. aeruginosa* was flushed from the microfluidic chambers before antibiotic treatment to test whether *P. aeruginosa* lysis products are responsible for the observed tolerance pattern. **(B)** Overall regrowth probability per chamber with *P. aeruginosa* present (yellow) or removed (grey). Each point represents one chamber; horizontal lines indicate medians. Statistical analysis: Welch's t-test; ns = not significant. **(C)** Regrowth probability as a function of distance from the porous wall. Lines show means; shaded regions indicate SD. The *P. aeruginosa*-present condition (yellow) represents the same experimental data as shown in Figures 1 and 2.

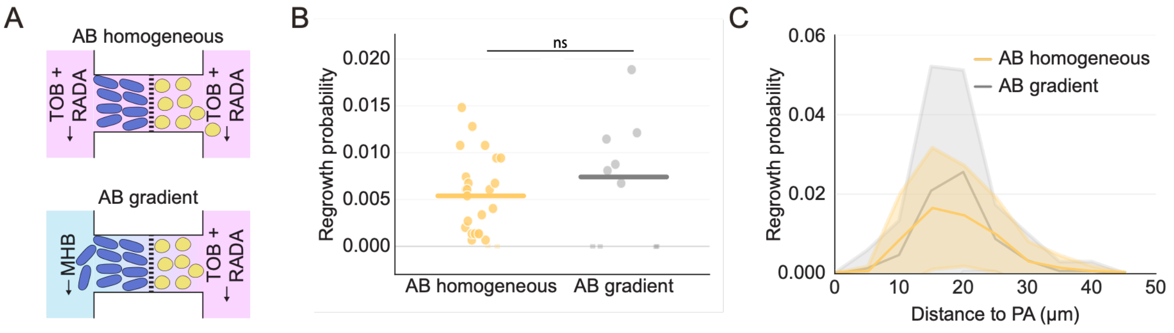


**Figure S9. Antibiotic gradient is not responsible for *S. aureus* tolerance pattern. (A)** To test whether tobramycin gradients are responsible for the observed tolerance pattern, tobramycin was delivered either homogeneously from both flow channels or as a gradient by delivery only from the *S. aureus*-side flow channel, resulting in the highest antibiotic concentration on the *S. aureus* side and the lowest concentration at the porous wall. **(B)** Overall regrowth probability per chamber under homogeneous (AB homogenous, yellow) or gradient (AB gradient, grey) antibiotic delivery conditions. Each point represents one chamber; horizontal lines indicate medians. Statistical analysis: Welch's t-test; ns = not significant. **(C)** Regrowth probability as a function of distance from *P. aeruginosa*. Lines show means; shaded regions indicate SD. The homogeneous antibiotic condition (yellow) represents the same experimental data as the *P. aeruginosa* wild-type condition shown in Figures 1 and 2.

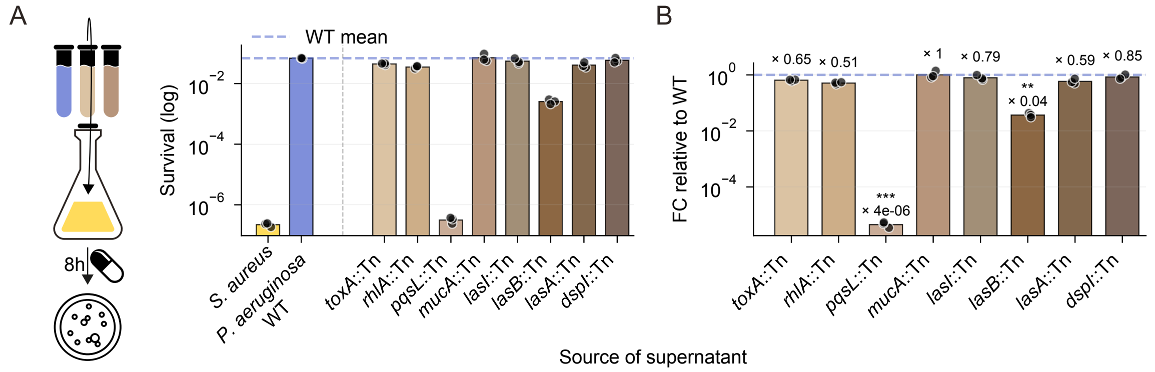


**Figure S10. Screening for *P. aeruginosa* factors affecting *S. aureus* tobramycin tolerance.** *S. aureus* was grown to mid-exponential phase in MHB medium and pre-treated with sterile supernatants from *P. aeruginosa* wild-type (WT) or transposon insertion mutants (::Tn) for 3 h. Subsequently, an aliquot was plated to determine baseline CFU count ($CFU_0$), and the remaining cells were treated with 58 µg/ml tobramycin for 8 h. All experiments were performed in biological triplicate. **(A)** Survival fraction ($CFU/CFU_0$) at 8h post-antibiotic treatment, shown on a $\log_{10}$ scale. **(B)** Fold change in survival relative to *P. aeruginosa* wild-type (WT) (dashed line). Bars represent geometric means; individual points show biological replicates. Statistical analysis: one-sample t-test on $\log_2$-transformed fold changes with Bonferroni correction. **$p < 0.01$, ***$p < 0.001$.

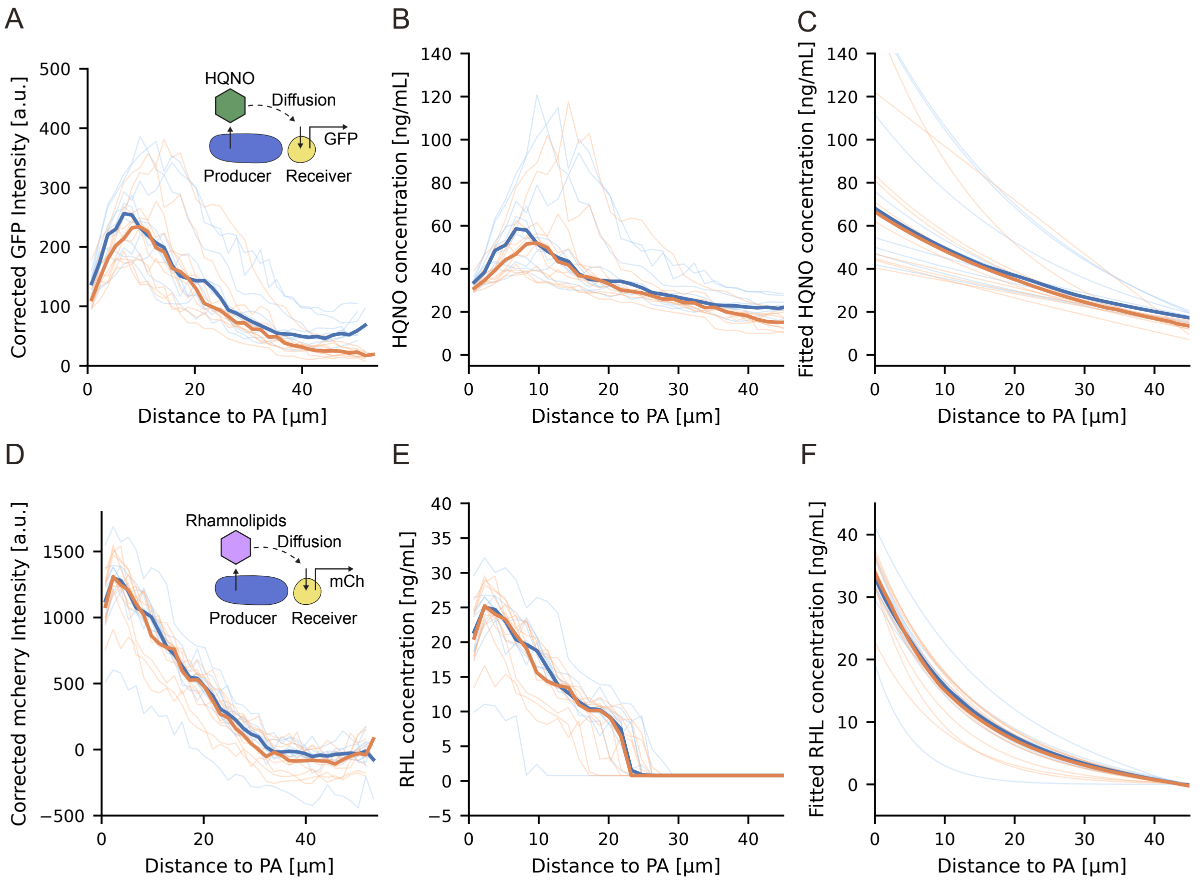


**Figure S11. Quantification of HQNO and rhamnolipid concentration gradients in microfluidic chambers.** Spatial gradients of *P. aeruginosa* secreted factors were measured in *S. aureus* using a transcriptional reporter (HQNO response) or propidium iodide staining (rhamnolipid-induced membrane permeabilization). **(A)** Background-corrected GFP fluorescence intensity (*pflB*-GFP transcriptional reporter) as a function of distance from *P. aeruginosa*. **(B)** HQNO concentration estimated from *pflB*-GFP intensity using calibration curves (Figure S12A–B). **(C)** Analytical solution of the reaction-diffusion model (Eq. 1, Methods) fitted to the HQNO concentrations in (B). Curves show the fitted profiles with inferred parameters. **(D)** Background-corrected propidium iodide fluorescence intensity as a function of distance from *P. aeruginosa*. **(E)** Rhamnolipid concentration estimated from propidium iodide intensity using calibration curves (Figure S12C–D). **(F)** Analytical solution of the reaction-diffusion model (Eq. 1, Methods) fitted to the rhamnolipid concentrations in (E). Curves show the fitted profiles with inferred parameters. Colors indicate independent microfluidic devices, thin lines represent individual chambers; thick lines show medians per device.

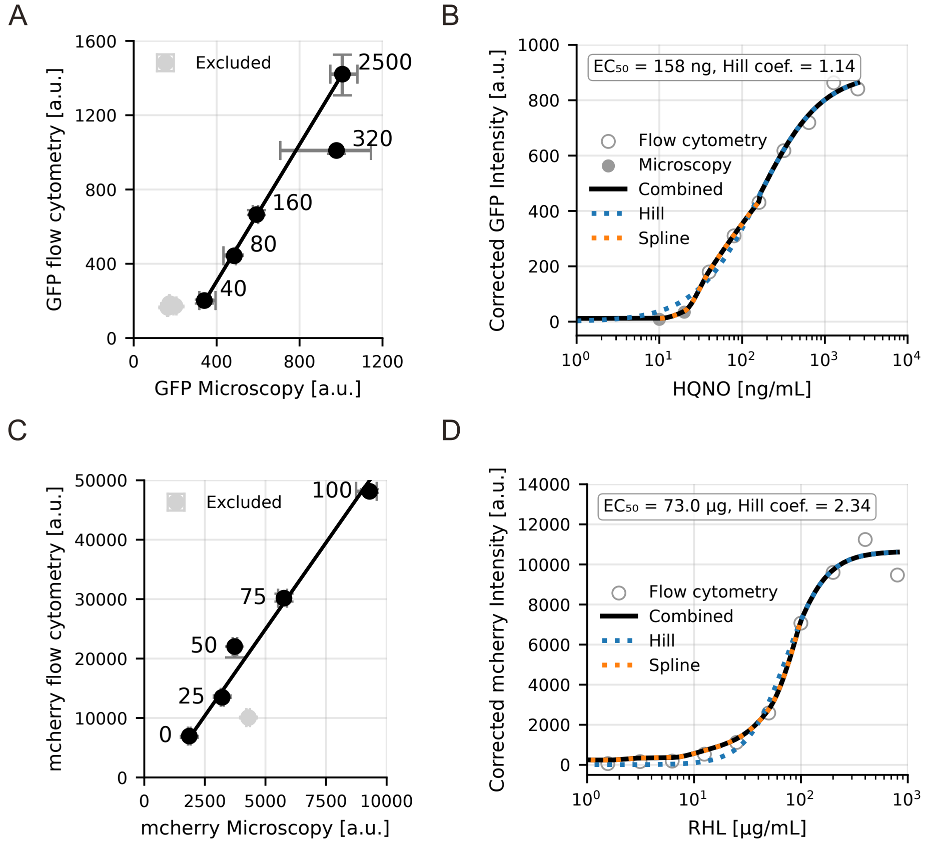


**Figure S12. Calibration curves for converting fluorescence intensity to HQNO and rhamnolipid concentrations. (A)** Correlation between GFP fluorescence (*pflB*-gfp) measured by flow cytometry and microscopy. Numbers indicate HQNO concentration (ng/mL); dots show median values and error bars indicate interquartile range (IQR); line shows result of ODR regression fit; light grey points were excluded from fit. **(B)** HQNO dose-response curve. Background-corrected GFP intensity in microscopy units plotted against HQNO concentration. Open circles: flow cytometry; filled circles: microscopy. Dashed blue line: Hill-curve fit ($EC_{50}$ = 158 ng/mL, Hill coefficient = 1.14); dashed orange line: PCHIP spline fit. Solid line: combined Hill-spline fit. **(C)** Correlation between propidium iodide fluorescence (mCherry channel) measured by flow cytometry and microscopy. Numbers indicate rhamnolipid concentration (µg/mL); light grey point was excluded from fit. **(D)** Rhamnolipid dose-response curve. Background-corrected propidium iodide intensity in microscopy units plotted against rhamnolipid concentration. Open circles: flow cytometry; filled circles: microscopy. Dashed blue line: Hill-curve fit ($EC_{50}$ = 73.0 µg/mL, Hill coefficient = 2.34); dashed orange line: PCHIP spline fit. Solid line: combined Hill-spline fit.

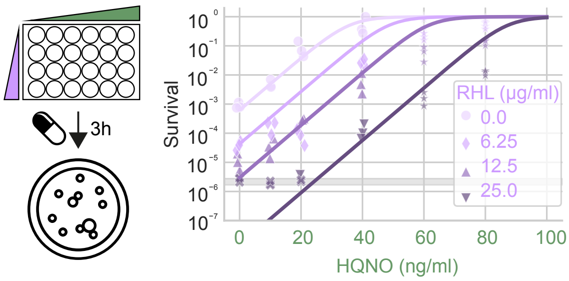


**Figure S13. Predicting *S. aureus* survival from HQNO and rhamnolipid concentrations.** Normalized survival probabilities of *S. aureus* as a function of HQNO concentration (x-axis) and rhamnolipid concentration (colors and symbols). *S. aureus* was grown to mid-exponential phase in MHB medium in 24-well plates, exposed to defined concentrations of HQNO and rhamnolipids for 3 h, and treated with 58 µg/ml tobramycin for 3 hours. Survival probabilities were determined using CFU counts and normalized by dividing by the highest observed level to account for growth during treatment initiation. Lines show predictions from a logistic regression model fitted to data with HQNO ≤ 60 ng/ml ($R^2 = 0.75$). Grey shaded area: limit of detection (LOD). Crosses: data points below LOD.

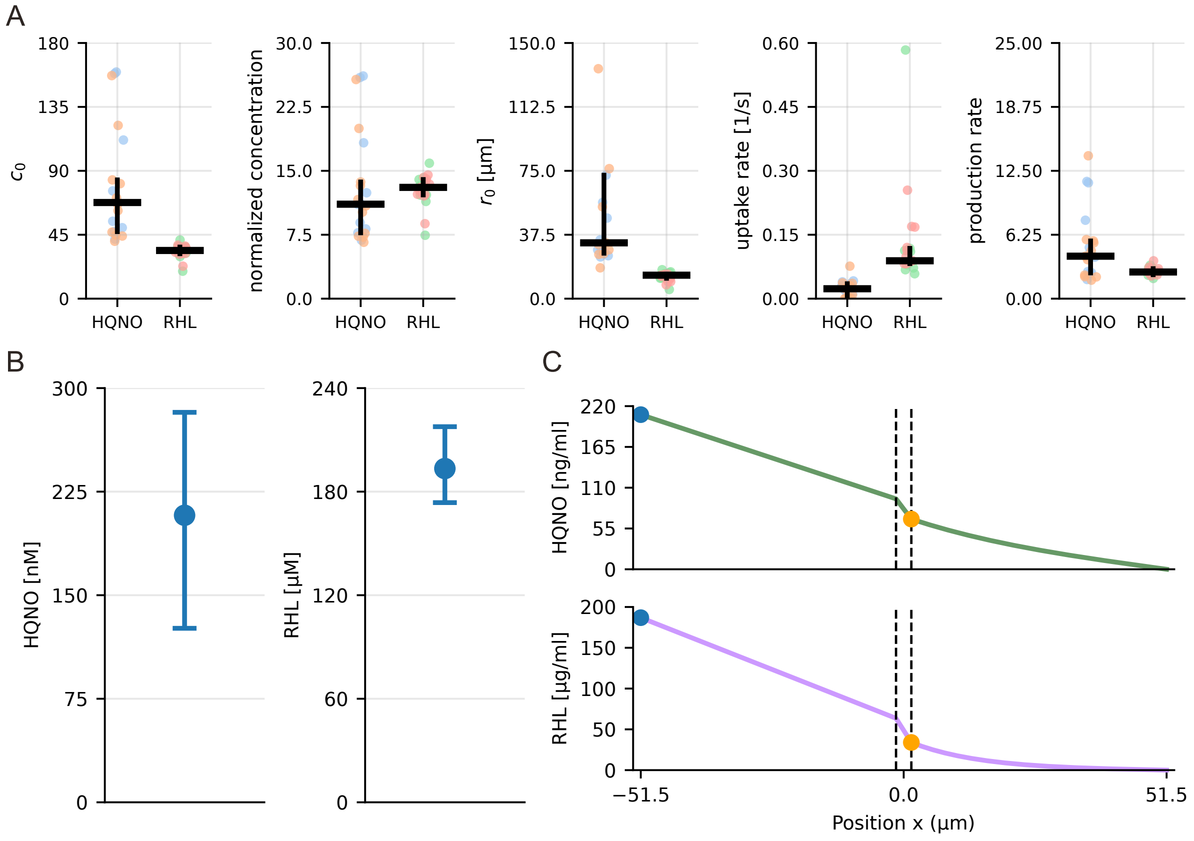


**Figure S14. Inferred reaction-diffusion model parameters. (A)** Parameter estimates from fitting the reaction-diffusion model to measured concentration gradients (Figure S11C, F). From left to right: maximum concentration, $c_0$, reached at the porous wall separating *S. aureus* from *P. aeruginosa*; normalized maximum concentration obtained by scaling $c_0$ by the logistic regression coefficient; interaction range, $r_0$; uptake rate, u; and production rate, p. Colors represent different microfluidic devices; points: individual chambers; lines: median with interquartile range. **(B)** Exogenous concentrations required in the flow channel on the *P. aeruginosa* side to reproduce the concentration gradients naturally produced by *P. aeruginosa*. **(C)** Predicted HQNO (top) and rhamnolipid (bottom) concentration profiles (median value). Dashed lines indicate porous wall region between *S. aureus* and *P. aeruginosa*; blue points mark exogeneous concentrations added to the flow channel; orange points mark maximum concentrations at the porous wall in the *S. aureus* chamber.

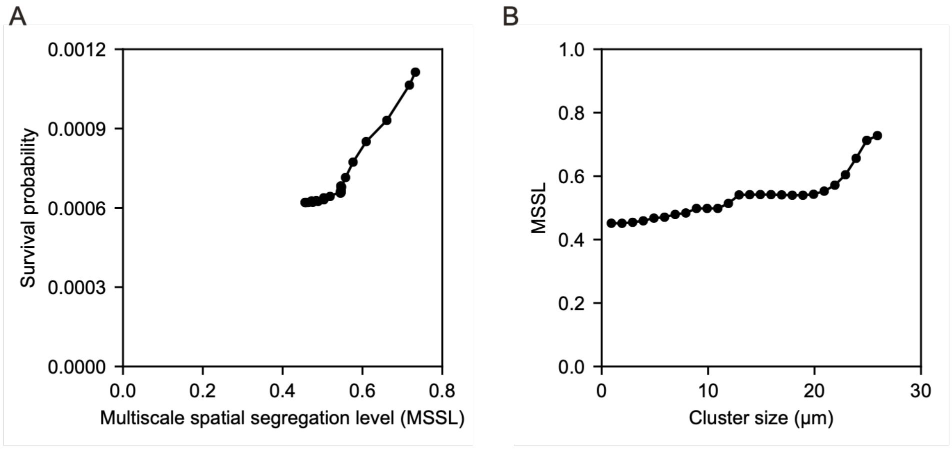


**Figure S15. Patch size of checkerboard pattern correlates with multiscale spatial segregation level.** Simulations were run using 2D models with checkerboard patterns of patch sizes ranging from 1 to 25 μm. **(A)** Average per-chamber survival probability as a function of multiscale spatial segregation level. Each point represents one checkerboard configuration. **(B)** Multiscale spatial segregation level as a function of patch size, showing positive correlation.

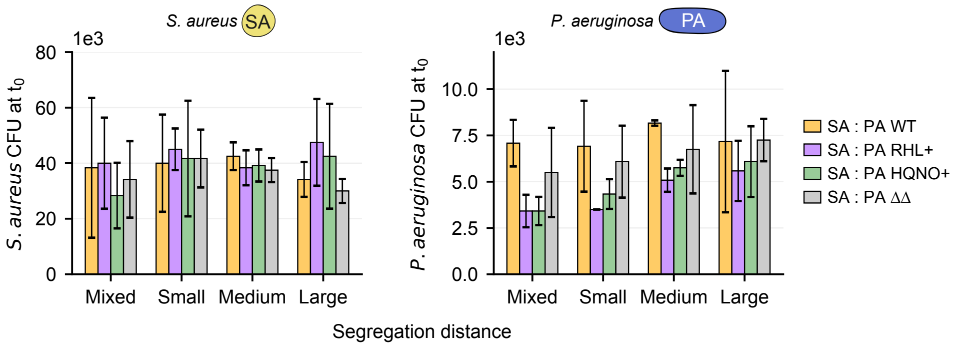


**Figure S16. Initial bacterial densities are comparable across checkerboard co-culture conditions.** *S. aureus* (left) and *P. aeruginosa* (right) $CFU_0$ before antibiotic treatment across spatial segregation patterns and *P. aeruginosa* strains (WT: wild-type; RHL+: rhamnolipid overproducer; HQNO+: HQNO overproducer; ΔΔ: *ΔpqsL ΔrhlA* double mutant). Bars: mean ± SEM. No significant differences were observed between strains within any spatial pattern. Statistical analysis: Kruskal-Wallis test, n = 3.

**Supplementary Movie 1. Survival of *S. aureus* in presence of *P. aeruginosa* wild type.** Microscopy image of *S. aureus* (right chamber) in presence of *P. aeruginosa* wild type (WT, left chamber, not shown) post tobramycin (58 µg/mL) and RADA (25 µM) removal, overlaying phase contrast (grey scale) with RADA fluorescence (magenta). Regrowing cells appear black, non-growing cells remain magenta. As cells regrow, they form clusters and push surrounding cells towards the flow channel on the right.

**Supplementary Movie 2. No regrowth in absence of *P. aeruginosa*.** Microscopy image of *S. aureus* in presence of *S. aureus* post tobramycin (58 µg/mL) and RADA (25 µM) removal, overlaying phase contrast (grey scale) with RADA fluorescence (magenta).

**Supplementary Movie 3. Survival of *S. aureus* in presence of *P. aeruginosa* rhamnolipid mutant.** Microscopy image of *S. aureus* (right chamber) in presence of *P. aeruginosa* rhamnolipid mutant (*ΔrhlA*, left chamber, not shown) post tobramycin (58 µg/mL) and RADA (25 µM) removal, overlaying phase contrast (grey scale) with RADA fluorescence (magenta). Regrowing cells appear black, non-growing cells remain magenta. As cells regrow, they form clusters and push surrounding cells towards the flow channel on the right.

**Table S1. List of strains and plasmids used.**

| Strains | Genotype | Description | Resistance | Reference |
|---|---|---|---|---|
| ***P. aeruginosa*** | | | | |
| SVP005 | Δ*pelA*-Δ*pslABC*-Δ*pilA*-Δ*fliC* | No motility | | [1] |
| SVP020 | pSEVA P*rhlA*-mcherry P*pqsL*-mNG in SVP005 background | HQNO and rhamnolipids promoter reporter | Km500 | This study |
| SVP023 | Δ*rhlA* KO mutant in SVP005 background | No rhamnolipids production | | This study |
| SVP025 | Δ*pqsL* KO mutant in SVP005 background | No HQNO production | | This study |
| SVP029 | Δ*rhlA* Δ*pqsL* KO mutant in SVP005 background | No rhamnolipids and HQNO production | | This study |
| SVP037 | pQFT-*rhlAB* in SVP005 background | Cumate-inducible vector expressing rhamnolipid synthesis genes | Carb100 | This study |
| SVP038 | pQFT-*pqsL* in SVP005 background | Cumate-inducible vector expressing HQNO synthesis genes | Carb100 | This study |
| SVP040 | pQFT-*rhlAB* in SVP023 background | Cumate-inducible vector expressing rhamnolipid synthesis genes in rhamnolipid KO mutant | Carb100 | This study |
| SVP041 | pQFT-*pqsL* in SVP029 background | Cumate-inducible vector expressing HQNO synthesis genes in HQNO KO mutant | Carb100 | This study |
| SVP046 | mCTX::mNeonGreen in SVP005 background | mNeonGreen tag | Oxytet100 | This study |
| SVP053 | pSEVA P*rhlA*-mcherry P*pqsL*-mNG in SVP005 background | HQNO and rhamnolipids promoter reporter | Cm30 | This study |
| SVP054 | pSEVA P*rhlA*-mcherry P*pqsL*-mNG in SVP029 background | HQNO and rhamnolipids promoter reporter | Cm30 | This study |
| SVP072 | mCTX::mNeonGreen in SVP029 background | mNeonGreen tag in rhamnolipid and hqno deficient mutant | Oxytet100 | This study |
| SVP075 | mCTX::mNeonGreen in SVP037 background | mNeonGreen tag in pQFT-*rhlAB* strain | Carb100 Oxytet100 | This study |
| SVP076 | mCTX::mNeonGreen in SVP038 background | mNeonGreen tag in pQFT-*pqsL* strain | Carb100 Oxytet100 | This study |

**Table S2—Continued. List of strains and plasmids used.**

| Strains | Genotype | Description | Resistance | Reference |
|---|---|---|---|---|
| ***S. aureus*** | | | | |
| SVS002 | BV18 MRSA USA300 | wild-type | | Bumann Lab, University of Basel |
| SVS021 | pHC48 in SVS002 background | RFP tag on pHC48 plasmid | Cm10 | This study |
| SVS021 | pHC48 pflB-gfp p4-mcherry in SVS002 background | pflB promoter reporter | Cm10 | This study |
| **Plasmids** | | | | |
| SVE002 | *E.coli* DH5$\alpha$ pex18Gm | suicide vector | Gm20 | Urs Jenal |
| SVE004 | *E. coli* top10 pex18Gm::*pqsL* | For constructing *pqsL* KO mutant | Gm20 | This study |
| SVE008 | *E.coli* DH5$\alpha$ pSEVA331 P*rhlA*-mcherry P*pqsL*-mNG | HQNO and rhamnolipids promoter reporter | Cm30 | This study |
| SVE013 | *E. coli* top10 pex18Gm::*rhlA* | For constructing *rhlA* KO mutant | Gm20 | This study |
| SVE017 | *E. coli* ec1000 pir+ $\Delta$dcm pHC48 *pflB*-gfp p4-mcherry | *pflB* promoter reporter | Amp100 | This study |
| SVE023 | *E. coli* top10 pQFT-*pqsL* | Cumate-inducible vector expressing HQNO synthesis genes | Amp100 | This study |
| SVE031 | *E. coli* top10 pQFT-*rhlAB* | Cumate-inducible vector expressing rhamnolipids synthesis genes | Amp100 | This study |

**Table S3. Primer sequences used.**

| Primer ID | Name | Target | Application | Sequence (5' -> 3') |
|---|---|---|---|---|
| svv001 | *rhlA*_dw_F | *rhlA* downstream region | pex18Gm::*rhlA* | GGAGGTGTGAAATGCGGCGCGAAAG TGGATTCCACGAGATGGCCATCGGC |
| svv002 | *rhlA*_dw_R | *rhlA* downstream region | pex18Gm::*rhlA* | ATTTCACACAGGAAACAGCTATGACCA TGATTACGAATTCATACTGCCGTCGAA CAGCGGG |
| svv003 | *rhlA*_up_F | *rhlA* upstream region | pex18Gm::*rhlA* | CCAGTCACGACGTTGTAAAACGACGG CCAGTGCCAAGCTTAGAAAAGCTTCG TCGACATCAACAGCG |
| svv004 | *rhlA*_up_R | *rhlA* upstream region | pex18Gm::*rhlA* | GCCGATGGCCATCTCGTGGAATCCAC TTTCGCGCCGCATTTCACACCTCC |
| svv005 | *rhlA*_seq_F | sequencing primer | sequencing pex18Gm::*rhlA* | TGGTCGATCCGAAGAACTTCGACG |
| svv006 | *rhlA*_seq_R | sequencing primer | sequencing pex18Gm::*rhlA* | TTCGTCGTCGAGCGGGGTCCC |
| svv007 | *pqsL*_dw_F | *pqsL* downstream region | pex18Gm::*pqsL* | CGGAGACTCATCCATGACGGACAACC ATCAGCCGGTGCGGTCGCCGGC |
| svv008 | *pqsL*_dw_R | *pqsL* downstream region | pex18Gm::*pqsL* | ATTTCACACAGGAAACAGCTATGACCA TGATTACGAATTCACCTCCAACCGCC CGGACTGG |
| svv009 | *pqsL*_up_R | *pqsL* upstream region | pex18Gm::*pqsL* | GCCGGCGACCGCACCGGCTGATGGT TGTCCGTCATGGATGAGTCTCCG |
| svv010 | *pqsL*_seq_F | sequencing primer | sequencing pex18Gm::*pqsL* | ACCTGGTGGCGCAACACGTGG |
| svv011 | *pqsL*_seq_R | sequencing primer | sequencing pex18Gm::*pqsL* | TGGCGCGGCTATTTCCCTCTTGGC |
| svv012 | *pqsL*_up_F | *pqsL* upstream region | pex18Gm::*pqsL* | CCAGTCACGACGTTGTAAAACGACGG CCAGTGCCAAGCTTAACGGCGCGTTC TGGAACATGGG |
| svv030 | pQFT_bla_F | introduce AmpR resistance | pQFT-Amp | TCGCCTTGCAGCACATCCCCCTTTCG CCAGCTGCGCGGAACCCCTATTTGTT TATTTTTCTAAATACATTC |
| svv031 | pQFT_bla_R | introduce AmpR resistance | pQFT-Amp | AGGTCAGGCTGGTGAGCGCCGCCAG TGAGCCTTGTTACCAATGCTTAATCAG TGAGGCACCTATCTCAGC |
| svv025 | *rhlA*_F | *rhlAB* gene amplification | pQFT-Amp *rhlAB* | AACAAACAGACAATCTGGTCTGTTTGT AACTAGTGCCTGTTCGAAAATTTTTGG GAGGTGTGAAATGC |
| svv038 | *pqsL*_F | *pqsL* gene amplification | pQFT-Amp *pqsL* | AACAAACAGACAATCTGGTCTGTTTGT AACTAGTTTCCGCCAGGAACGACACG GAGACTCATCC |
| svv039 | *rhlB*_R | *rhlAB* gene amplification | pQFT-Amp *rhlAB* | TAGTCCGGATCCCAATTGGAGCTCGG TACCTCAGGACGCAGCCTTCAGCCAT CGAGCATCC |
| svv040 | *pqsL*_R | *pqsL* gene amplification | pQFT-Amp *pqsL* | TAGTCCGGATCCCAATTGGAGCTCGG TACCTCAGCCGAGCGGCGCCGGCGA CC |
| svv056 | pQFT_seq_F | sequencing primer | sequencing pQFT-Amp | TACATATGTCAATGTACCGG |
| svv057 | pQFT_seq_F | sequencing primer | sequencing pQFT-Amp | TGTAAAACGACGGCCAG |

| | | | | |
|---|---|---|---|---|
| svv112 | pflB_F | *pflB* gene amplification | pHC48 *pflB*-gfp p4-mcherry | CGACTCTAGAGGATCCGCTAGCCTGTCAGACCAAGTTAAAAAGCGCA |
| svv013 | pflB_R | *pflB* gene amplification | pHC48 *pflB*-gfp p4-mcherry | ACAGCTATGACATGATTACGAATTCATAATGCCGACTGTACTTTTTACATCAC |
| svv070 | pflB_seq_R | sequencing primer | sequencing pHC48 *pflB*-gfp p4-mcherry | ACCTTCGGGTGGGCCTTT |
| svv122 | pflB_seq_F | sequencing primer | sequencing pHC48 *pflB*-gfp p4-mcherry | ACTGGGCAGTGTCTTAAAAAATCG |
| svv158 | mNeon_F | introduce mNeon into pSEVA | pSEVA P*rhlA*-mcherry P*pqsL*-mNG | ATTCGAGCTCGGTACCCGGGGATCCTGAAGCGAAGGAGGTTACGACGAAATATGGTGAGCAAAGGTGAAGAGG |
| svv159 | mNeon_R | introduce mNeon into pSEVA | pSEVA P*rhlA*-mcherry P*pqsL*-mNG | TCACGACGCGGCCGCAAGCTTTATTTGTACAGCTCATCC |
| svv160 | PpqsL_F | amplify pqsL promoter | pSEVA P*rhlA*-mcherry P*pqsL*-mNG | GCCTAGGCCGCGGCCGCGCGAATTCgatcgtcaccgtcaactgc |
| svv161 | PpqsL_R | amplify pqsL promoter | pSEVA P*rhlA*-mcherry P*pqsL*-mNG | GGATCCCCGGGTACCGAGCTCggatgagtctccgtgtcgttcc |
| svv162 | PrhlA_F | amplify PrhlA promoter | pSEVA P*rhlA*-mcherry P*pqsL*-mNG | GAGCTGTACAAATAAAGCTTATTTGTCCTACTCAGGAGAGCGTTCACCGACAAACAACAGATAAAACGAAAGGCCCAGTCTTTCGACTGAGCCTTTCGTTTTATTTGagatctacgccaatgaaggcg |
| svv163 | PrhlA_R | amplify PrhlA promoter | pSEVA P*rhlA*-mcherry P*pqsL*-mNG | TGTCTCTTAACGTAACCGTAGGATCCCCGGGTACCGAGCTC |
| svv164 | mCherry_F | introduce mCherry2 into pSEVA | pSEVA P*rhlA*-mcherry P*pqsL*-mNG | TACGGTTACGTTAAGAGACATTTATGG |
| svv165 | mCherry_R | introduce mCherry2 into pSEVA | pSEVA P*rhlA*-mcherry P*pqsL*-mNG | AACAGGAGTCCAAGACTAGTTTACTTGTACAGCTCGTCCATGCC |

**Table S3. Model parameters.** Parameters used in the reaction-diffusion model. † Interaction range refers to the characteristic length-scale of reaction-diffusion process. ‡ Coefficients of regression analysis relating *S. aureus* survival probability in batch culture to HQNO and RHL concentrations: $P(\text{survival}) = \sigma(a + b \cdot [\text{HQNO}] + c \cdot [\text{RHL}])$, where $\sigma(z) = \frac{1}{1+e^{-z}}$ is the logistic function. RHL: rhamnolipids. Values in brackets indicate interquartile range.

| Symbol | Description | Value | Source |
|---|---|---|---|
| $\mathrm{D_{HQNO}}$ | Diffusion constant HQNO in empty space | 27 μm²/s | [2] |
| $\mathrm{D_{RHL}}$ | Diffusion constant RHL in empty space | 40 μm²/s | Estimated using Stokes-Einstein-Sutherland equation |
| $\rho$ | Cell density (occupied volume fraction) | 0.46 | Measured |
| $\mathrm{r_{0,HQNO}}$ | Interaction range HQNO† | 32.8 (26.7 – 72.5) μm | Fitted to data |
| $\mathrm{r_{0,RHL}}$ | Interaction range RHL† | 13.8 (12.0 – 14.3) μm | Fitted to data |
| $\mathrm{c_{0,HQNO}}$ | HQNO concentration at porous wall | 67.7 (47.2 – 83.6) ng/ml | Fitted to data |
| $\mathrm{c_{0,RHL}}$ | RHL concentration at porous wall | 33.9 (31.6 – 36.3) μg/ml | Fitted to data |
| $\mathrm{u_{HQNO}}$ | Uptake rate HQNO in *S. aureus* | 0.023 (0.0048 – 0.035) 1/s | Calculated using $u = D/r_0^2$ |
| $\mathrm{u_{RHL}}$ | Uptake rate RHL in *S. aureus* | 0.089 (0.082 – 0.12) 1/s | Calculated using $u = D/r_0^2$ |
| $\mathrm{p_{HQNO}}$ | Production rate HQNO in *P. aeruginosa* | 4.2 (2.5 – 5.6) ng/ml/s | Calculated using Eq. 2 |
| $\mathrm{p_{RHL}}$ | Production rate RHL in *P. aeruginosa* | 2.6 (2.3 – 2.9) μg/ml/s | Calculated using Eq. 2 |
| $\alpha$ | Porosity of wall (width of gaps relative to chamber width) | 0.25 | Measured |
| d | Thickness porous wall | 3 μm | Measured |
| L | Chamber length (distance from porous wall to flow channel) | 50 μm | Measured |
| a | Regression intercept‡ | -7.10 | Fitted to data |
| b | Regression coefficient HQNO‡ | 0.164 ml/ng | Fitted to data |
| c | Regression coefficient RHL‡ | -0.385 ml/ μg | Fitted to data |
